# Prenatal immune stress induces a prolonged blunting of microglia activation that impacts striatal connectivity

**DOI:** 10.1101/2021.12.27.473694

**Authors:** Lindsay N. Hayes, Kyongman An, Elisa Carloni, Fangze Li, Elizabeth Vincent, Manish Paranjpe, Gül Dölen, Loyal A. Goff, Adriana Ramos, Shin-ichi Kano, Akira Sawa

## Abstract

Recent studies suggested that microglia, the primary brain immune cells, can affect circuit connectivity and neuronal function^1–3^. Microglia infiltrate the neuroepithelium early in embryonic development and are maintained in the brain throughout adulthood^4,5^. Several maternal environmental factors, such as aberrant microbiome, immune activation, and poor nutrition, can influence prenatal brain development^6–8^. Nevertheless, it is unknown how changes in the prenatal environment instruct the developmental trajectory of infiltrating microglia, which in turn affect brain development and function. Here we show that after maternal immune activation (MIA) microglia from the offspring have a long-lived decrease in immune reactivity (blunting) across the developmental trajectory. The blunted immune response was concomitant with changes in the chromatin accessibility and reduced transcription factor occupancy of the open chromatin. Single cell RNA sequencing revealed that MIA does not induce a distinct subpopulation but rather decreases the contribution to inflammatory microglia states. Prenatal replacement of MIA microglia with physiological infiltration of naïve microglia ameliorated the immune blunting and restored a decrease in presynaptic vesicle release probability onto dopamine receptor type-two medium spiny neurons, indicating that aberrantly formed microglia due to an adverse prenatal environment impacts the long-term microglia reactivity and proper striatal circuit development.

## Main

Microglia, the primary immune cell in the brain, have many functions across the lifespan. Although their primary function of rapid immune activation is to protect the brain from harm, dysfunction of the microglia immune activation can result in poor outcomes for the surrounding neurons and glia^9,10^. Since microglia are long-lived cells with early brain entry, aberrant microglia arriving in early development are likely to play a key role in modulating these cell-cell interactions across the trajectory of health and disease. Several studies showed an impact of prenatal stress on brain development and function in later life^11–14^. Nevertheless, the role and mechanism of microglia in the long-term changes elicited by prenatal stress are unknown.

### Blunted Microglia Reactivity in Adult MIA Offspring

To address this knowledge gap, we tested whether a prenatal immune stressor (maternal immune activation, MIA) influenced microglia functions including immune activation. To generate MIA offspring, we delivered an immune activator (polyinosinic-polycytidylic acid, PIC) to pregnant dams at embryonic day (E)9.5 to target the first wave of microglia infiltrating the neuroepithelium^4^. To study how MIA influences the offspring immune response, we injected either a pro-inflammatory stimulus, lipopolysaccharide (LPS), or saline (SAL) to adult offspring *in vivo*, isolated microglia, and analyzed the gene expression profiles (**Ext Data Fig. 1**). First, we compared the gene expression between microglia from MIA offspring (MIA microglia) and those from control offspring (CON microglia). We observed only 7 differentially expressed genes (DEGs) between MIA and CON microglia after SAL treatment (**Fig. 1a, Ext Data Fig. 2**). In contrast, we identified 401 DEGs between MIA and CON microglia after LPS treatment (**Fig. 1a, Ext Data Fig. 2**). Interestingly, the majority of the DEGs (76%) were downregulated in MIA compared with CON microglia (**Fig. 1a**). Furthermore, the DEGs were significantly enriched for pathways including downregulation of many immune response pathways (**Fig. 1b**). Altogether, adult immune reactivity is markedly reduced in MIA microglia, although the baseline (SAL) gene expression is similar between MIA and CON microglia.

**Figure 1:**
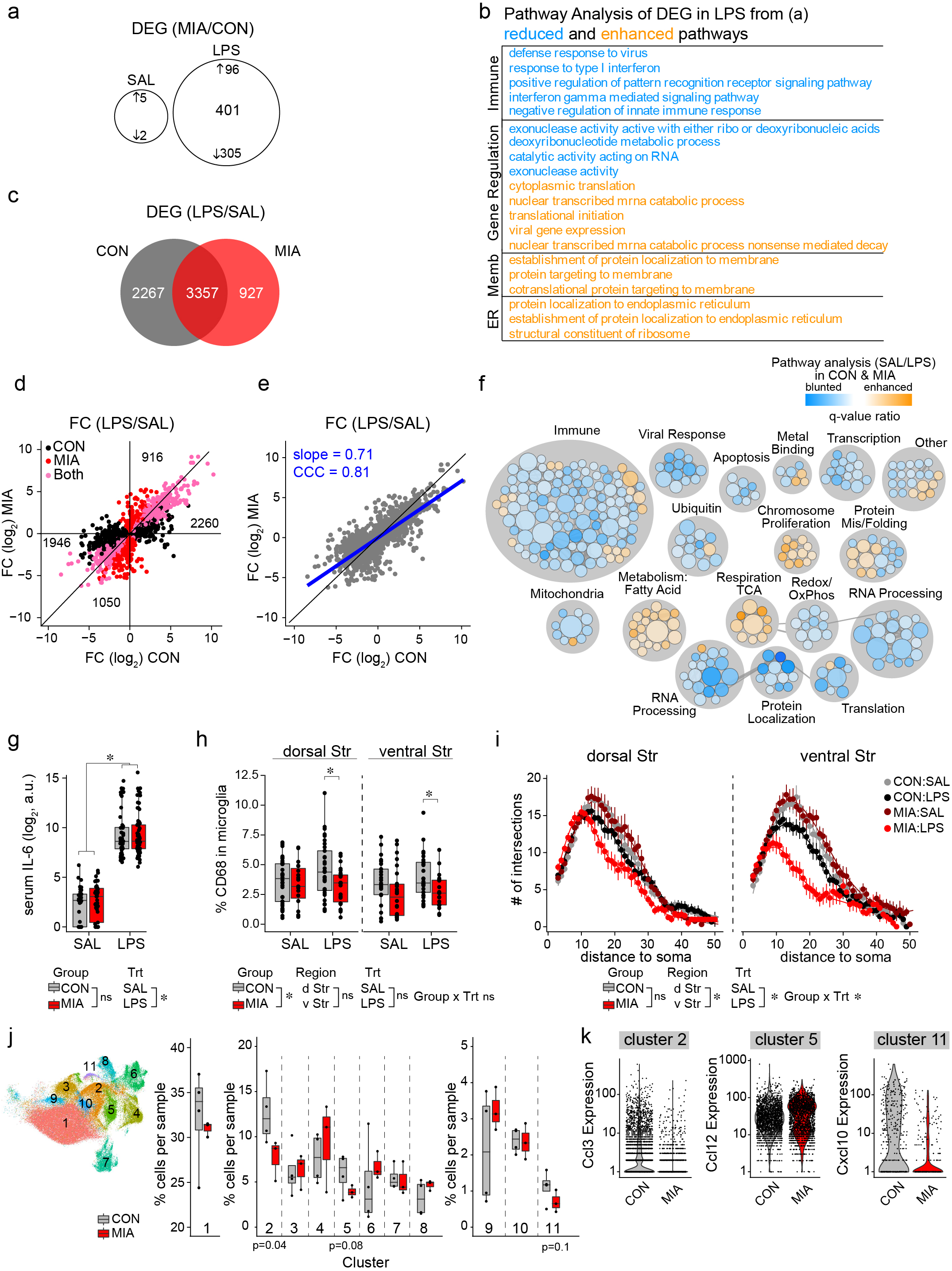
Diminished microglia immune response after MIA. **a,** Venn diagram of differentially expressed genes (DEGs, q <0.05) between maternal immune activation (MIA) and control (CON) microglia after acute saline (SAL) or lipopolysaccharide (LPS) treatment. **b,** Gene set enrichment analysis of DEGs in (a) between MIA and CON microglia after acute LPS treatment *in vivo* listing the top 20 gene ontology pathways with blue indicating reduction in MIA microglia and orange indicating enhancement in MIA microglia. **c**, Venn diagram of DEGs (q <0.05) between SAL and LPS in CON microglia compared with MIA microglia. **d-e,** Plot (d) and linear regression (e) of the fold changes (FC) for the LPS-induced DEGs in both CON and MIA microglia (pink), only CON microglia (black) or only MIA microglia (red). Numbers indicate the total number of LPS response genes in each section. The black line indicates the y=x line and the blue line is the linear fit line (slope = 0.71, concordance coefficient, CCC = 0.81). **f,** Comparison of the pre-ranked gene set enrichment analysis for LPS response genes between the CON and MIA microglia. Each circle represents a gene ontology term with the color indicating the magnitude of the q-value ratio (q <0.05). Terms were selected with a log_2_ q-value ratio ≥1. Blue is for a blunted response to LPS, and orange is for an enhanced response to LPS in MIA microglia. The grey lines represent the similarity coefficient (thickness) or the amount of genes the two pathways have in common. **g**, Serum interleukin-6 (IL-6) from CON and MIA offspring after SAL or LPS treatment. n are individual animals. Data are shown as median ± interquartile range and analyzed using a linear mixed effect model which showed an LPS effect, but no MIA effect * p<0.05, a.u. = arbitrary units. **h**, Percent CD68 content inside Iba1+ microglia in the dorsal and ventral striatum (Str). n are individual cells. Data are shown as median ± interquartile range and analyzed using a 3-way ANOVA (MIA, LPS, region). There is a significant main effect of MIA with a significant emmeans post-hoc test for MIA microglia versus CON microglia after LPS treatment of both brain regions * p<0.05. **i**, Morphological analysis of Iba1+ microglia using Sholl analysis. n are individual cells. Data are shown as mean ± s.e.m and analyzed using a linear mixed effect model which showed a significant effect of treatment (Trt), brain region (Region), and interaction between MIA group and LPS treatment (Group x Trt). * p<0.05. **j,** Uniform Manifold Approximation and Projection (UMAP) of ∼120,000 microglia grouped into 11 clusters. Differential abundance analysis quantifying the percent of cells from each sample in each cluster. n are individual animals. Data are shown as median ± interquartile range and analyzed using a t-test. **k,** Gene expression (log_10_) for the distinct marker genes for clusters 2, 5, and 11 from CON and MIA microglia. n are individual cells.

To address the decreased MIA microglia immune reactivity directly, we compared the microglia gene expression between LPS and SAL treatment (LPS response genes). In CON microglia, 5624 genes were differentially expressed between LPS and SAL treatment, whereas in MIA microglia fewer genes (4284) were differentially expressed between these two (**Fig. 1c, Ext Data Fig. 2**). Specifically, 71% of the LPS-induced genes (2260) showed a diminished induction (decreased gene expression) in MIA microglia compared with CON microglia (**Fig. 1d, Ext Data Fig. 2**). Similarly, 65% of the LPS-suppressed genes (1946) showed a diminished suppression (increased gene expression) in MIA compared with CON microglia (**Fig. 1d, Ext Data Fig. 2**). Together these results indicate a blunted immune response in MIA microglia compared to CON microglia, with a blunted gene defined as any gene with the LPS response in MIA microglia that has a smaller absolute fold change than the LPS response in CON microglia. We also demonstrated the immune blunting by fitting a regression line to the fold change values comparing the LPS response between CON and MIA microglia with a slope of ∼ 0.7 and a concordance coefficient of ∼ 0.8 both less than 1 (**Fig. 1e, Ext Data Fig. 2**). These data are in contrast to an expectation of a hyperactive immune response in MIA that was commonly believed^15–17^.

To functionally characterize this surprising result, we compared two independent gene set enrichment analyses to identify specific molecular pathways that were differentially enriched between MIA and CON microglia in the context of their response to *in vivo* LPS treatment in adulthood. We found a strong decrease in the innate immune response pathways including interferon signaling in MIA microglia (**Fig. 1f**). In addition, we found an enhancement of the tricarboxylic acid (TCA) cycle and fatty acid metabolism in parallel with a reduction in oxidative phosphorylation, redox pathways, electron transport chain, and respiratory pathways in MIA microglia (**Fig. 1f**). The imbalance of the mitochondrial pathways seen in MIA microglia suggests an impaired immunometabolic response. Altogether, these results emphasize broad functional adaptations that are implemented in microglia in response to a change in the maternal microenvironment.

We next addressed whether there is a regional specificity of the blunted LPS response in adult MIA microglia. We found that MIA microglia from the frontal cortex and striatum both showed a blunted immune response in comparison with CON microglia, but the striatum showed a more robust effect than the frontal cortex (**Ext Data Fig. 3a-b**). To determine the specificity of the immune blunting to MIA microglia, rather than a bias in the data, we compared the LPS response between microglia from the frontal cortex and those from the striatum (mixing both CON and MIA together) and found a more concordant immune response (slope = 0.9, CCC = 0.9) indicating the blunted immune response is specific to MIA microglia (**Ext Data Fig. 3c-e**). In addition, the peripheral immune response to LPS treatment between MIA and CON offspring was not different as measured by serum interleukin-6 (IL-6) indicating that the peripheral immune response was normal (**Fig. 1g**). Altogether, the diminished immune responsiveness is a specific feature associated with MIA microglia in the brain.

Next, we functionally validated these *in vivo* observations in adult offspring using histology (**Fig. 1h-i, Ext Data Fig. 4**). We measured the density, morphology, and lysosomal content of adult microglia in the dorsal and ventral striatum (**Fig. 1h-i, Ext Data Fig. 4**) because the striatal microglia showed a stronger immune phenotype (**Ext Data Fig. 3a-b**). Consistent with a blunted immune reactivity, we found that MIA microglia had less Cd68 lysosomes than CON microglia after *in vivo* LPS treatment (**Fig. 1h, Ext Data Fig. 4b-d**). Furthermore, MIA microglia were significantly smaller than CON microglia after LPS, although morphologic differences were negligible after SAL treatment, indicating an impaired responsiveness in MIA microglia (**Fig. 1i, Ext Data Fig. 4a**). The microglia density was unchanged upon both SAL and LPS treatments (**Ext Data Fig. 4e**). Altogether these data support a blunted immune response phenotype in MIA microglia by histology.

Several single cell RNA sequencing studies identified unique microglia subpopulations, particularly in neurodegenerative conditions^18–22^. Therefore, we evaluated whether the blunted immune response phenotype occurs in a distinct subset of microglia. Among all the LPS treated cells, we identified 11 distinct clusters of microglia, but no distinct cluster for MIA microglia compared with CON microglia (**Fig. 1j-k, Ext Data Fig 5a-c**). Next, we performed an abundance assay to quantify the proportion of cells from each sample to each cluster. We found that MIA had a reduced abundance in the proinflammatory cluster 2 (*Ccl3+*), and a decreasing trend in clusters 5 (*Ccl12+*) and 11 (interferon cluster, *Ifit2+ and Cxcl10+*) (**Fig. 1j-k, Ext Data Fig 5**). These data indicate that MIA affects some proinflammatory clusters, consistent with the idea of a blunted immune phenotype in MIA microglia observed in the bulk RNA sequencing analysis.

### Blunted Microglia Reactivity in Development after MIA

We next addressed the immune responsiveness of microglia in the developmental course after MIA, by extracting microglia from the striatum and cortex of both MIA and CON offspring and culturing the primary microglia *in vitro*. This allows us to characterize immune responsiveness of embryonic microglia and to obtain complementary findings with the *in vivo* data. We stimulated the extracted microglia with LPS *in vitro* and quantified the secretion of cytokines IL-6 and TNFα from the microglia into the culture media. Consistent with the *in vivo* data, we found that adult microglia from the striatum of MIA offspring showed a reduction in the secreted proinflammatory cytokines compared with CON microglia (**Fig. 2a**). Importantly, a similar diminished immune response was observed at E18 in MIA compared with CON microglia from the striatum, indicating a long-lived immune blunting phenotype (**Fig. 2a**). In cortical microglia, MIA showed a mild reduction in adulthood and no significant change at E18 in the immune response, indicating the cortex has a delayed and milder immune blunting phenotype potentially due to the later maturation of cortical development in comparison with the striatum (**Fig. 2b**). These *in vitro* data suggest that a prenatal immune stressor induced a long-lived blunting in MIA microglia activation to a subsequent immune stimulus, consistent with the blunted response of microglia to *in vivo* stimulation in adulthood.

**Figure 2:**
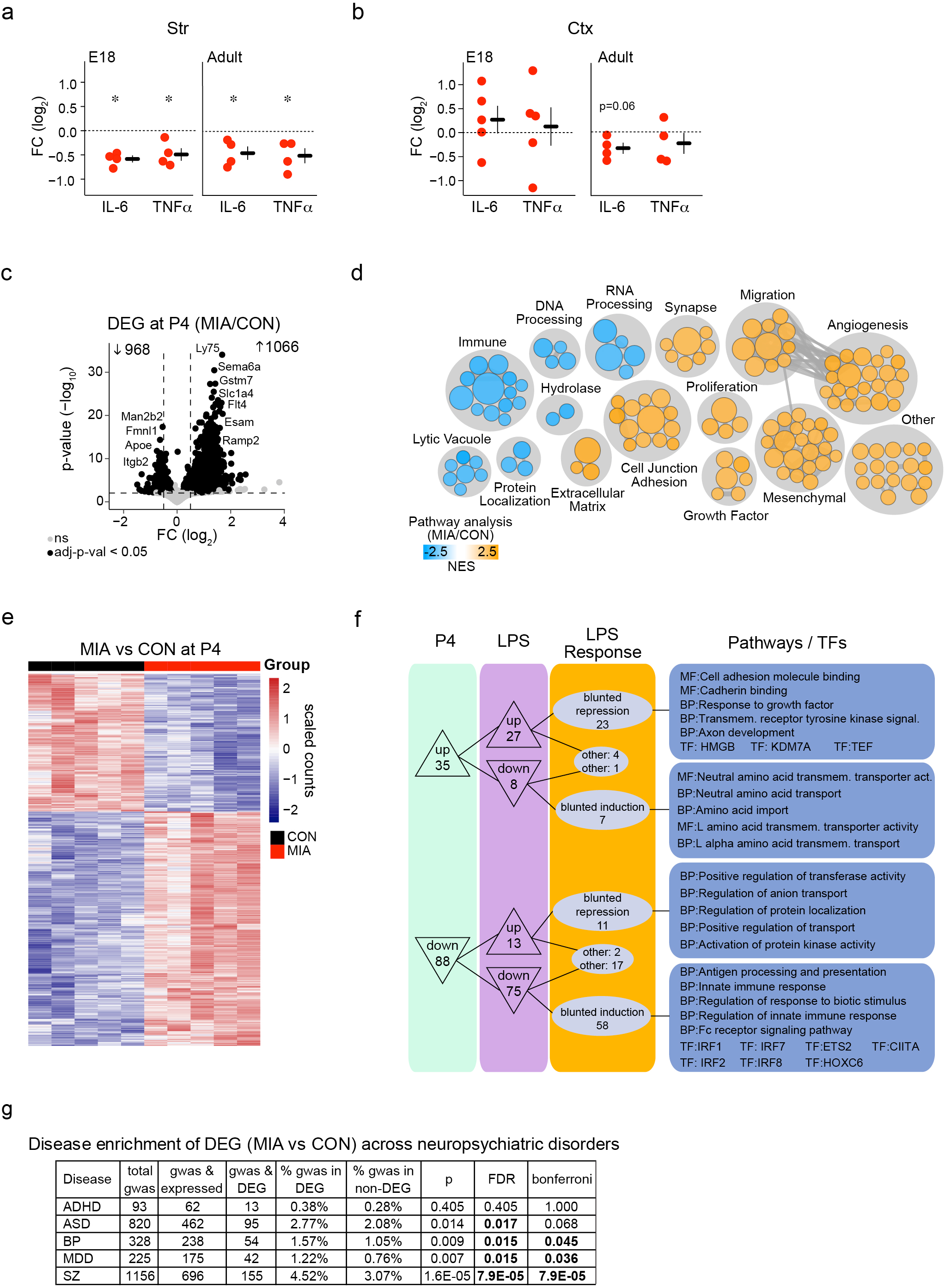
Diminished microglia reactivity in development after MIA. **a-b**, Quantification of secreted IL-6 and tumor necrosis factor alpha (TNFα) from LPS-stimulated microglia *in vitro* at embryonic day (E)18 and adulthood from the whole Str (a) and cortex (Ctx) (b). n are individual culture experiments. Data are shown as mean ± s.e.m. with each experiment normalized to CON and analyzed using a one-sample t-test. *p<0.05. **c,** Volcano plot of DEGs between MIA and CON microglia from the whole neonatal brain (black, q <0.05). **d**, Gene set enrichment analysis of neonatal DEGs between MIA and CON microglia with blue pathways diminished in MIA microglia and orange pathways enhanced in MIA microglia. **e**, Heatmap of neonatal DEGs between MIA and CON microglia. **f**, 123 overlapping DEGs (q < 0.1) between MIA and CON from neonatal microglia and LPS-treated adult microglia along with enriched pathways [molecular function (MF) and biological process (BP)] and transcription factor (TF) regulators. **g**, Enrichment of genome wide association study (GWAS) risk genes for neuropsychiatric disorders among DEGs between MIA and CON microglia using a chi-square test. Attention deficit hyperactivity disorder (ADHD), autism spectrum disorder (ASD), bipolar disorder (BP), major depressive disorder (MDD), and schizophrenia (SZ), p = p-value, FDR = Benjamini-Hochberg corrected p-value, bonferroni = Bonferroni corrected p-value.

The neonatal brain contains endogenous stressors that can induce microglia to become reactive^21,23–25^. Thus, by directly examining *in vivo* microglia from freshly extracted neonatal brain, we can evaluate MIA microglia responsivity to intrinsic activators. We found 2034 DEGs between neonatal MIA and CON microglia (**Fig. 2c-e**). Similar to LPS-activated microglia in adulthood (**Fig. 1b, f**), these DEGs between neonatal MIA and CON microglia showed a reduction in pathways associated with the immune response, phagocytosis, RNA processing, and protein localization (**Fig. 2d**). Moreover, 123 of these DEGs were commonly changed (FDR <0.1) in neonatal (intrinsically activated) microglia and adult LPS-treated microglia (**Fig. 2f**). Most of the genes were downregulated in neonatal microglia (88 genes) and in adult LPS-treated microglia (75 genes) (**Fig. 2f**). These genes were functionally related to the innate immune response and were enriched for target genes of the interferon pathway (Irf1, Irf2, Irf7, Irf8)^26^ (**Fig. 2f**). Altogether, MIA results in a diminished microglial reactivity to both intrinsic activation in the neonatal stage and exogenous activation by LPS in adulthood, with an impairment in overlapping functional pathways.

We also determined whether the DEGs between MIA and CON across datasets were enriched for disease-associated risk genes by examining datasets from genome-wide association studies (GWASs)^27^. We found DEGs between MIA and CON were enriched for schizophrenia risk genes more than risk genes for autism spectrum disorder, attention deficit hyperactivity disorder, bipolar disorder, and major depression disorder (**Fig. 2g**). Schizophrenia is a syndrome of neurodevelopmental origin with symptom onset after adolescence in contrast to child-onset developmental conditions such as autism spectrum disorder and attention deficit hyperactivity disorder or adult-onset conditions such as bipolar disorder and major depressive disorder^28,29^. As a result, the contribution of immune mechanisms to autism may precede schizophrenia across the developmental trajectory. Thus, the pathological involvement of microglia throughout the developmental trajectory may be important for schizophrenia. In summary, these data provide evidence for a common underlying pathophysiology between environmental stressors (i.e., MIA) and genetic risk factors on microglia biology in disease.

### Epigenetic Mechanism of MIA Microglia Immune Blunting

We looked for a mechanism for how MIA could induce the blunted immune phenotype in adulthood. We hypothesized that such a long-term impairment might be associated with an epigenetic change that interfered with transcriptional regulation of immune activation. To test this idea, we performed ATAC (Assay for Transposase Accessible Chromatin) sequencing of adult MIA and CON microglia after LPS treatment. Surprisingly, we found that 97% of the differentially accessible regions (DARs) were more open in MIA compared with CON microglia (**Fig. 3a**). After multiple comparison correction, we identified 113 more open DARs with approximately a quarter near transcriptional start sites (**Fig. 3b**). We used the Genomic Regions Enrichment of Annotations Tool (GREAT, version 4.0.4) to identify enriched pathways for the open chromatin regions including the aforementioned immune regulation, lysosomal, and protein modification pathways (**Fig. 3c**). However, a remaining question is how a more open chromatin structure could lead to a blunted transcriptional program.

**Figure 3:**
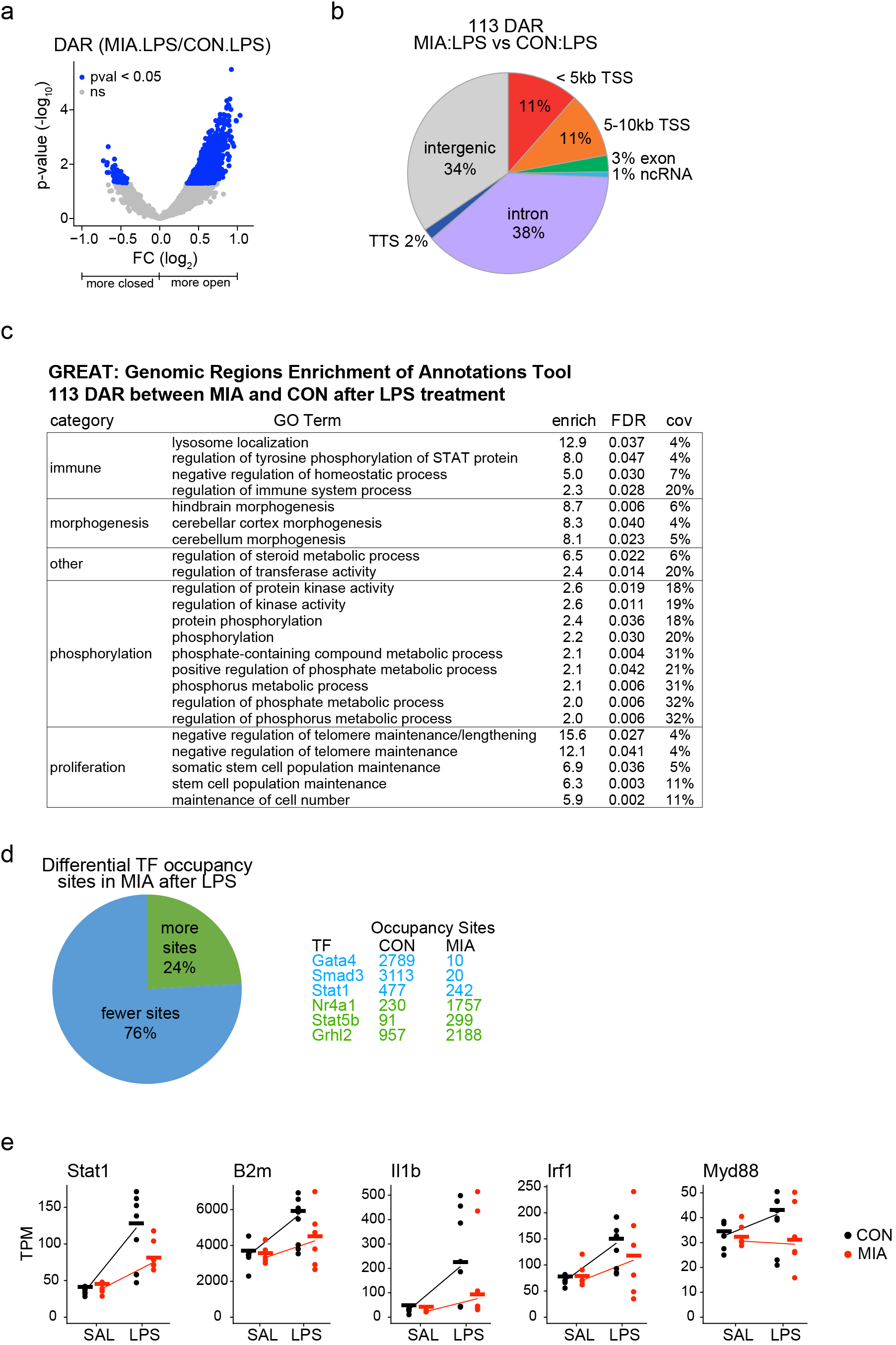
Epigenetic reprogramming and transcriptional regulation of MIA microglia. **a,** Volcano plot of differentially accessible regions (DARs) of assay for transposases accessible chromatin (ATAC) sequencing between MIA and CON microglia from the whole Str after LPS treatment. (blue, p <0.05). A positive FC indicates more open chromatin in MIA microglia, and a negative FC indicates more closed chromatin in MIA microglia. **b**, Pie chart of the spatial location of 113 DARs after permutation test (adjusted p <0.05) with the location respective to the transcriptional start site (TSS), exon, non-coding RNA (ncRNA), intron, intergenic, and transcriptional termination site (TTS). **c**, Pathway analysis for the 113 DARs between MIA and CON microglia after LPS treatment. 23 enriched gene ontology (GO) pathways with FDR < 0.05. Enrichment score (enrich) and coverage (cov). **d**, Pie chart of TF occupancy sites in the open chromatin regions of MIA and CON microglia after LPS treatment. TFs were included with more than 250 total sites and ≥ 0.5 log_2_ FC. Right table indicates the top 3 more and less occupied TFs with the number of occupancy sites in CON and MIA microglia after LPS. **e**, Gene expression (transcripts per million, TPM) of CON and MIA microglia after SAL or LPS treatment for *Stat1* and Stat1 target genes (*B2m*, *Il1b*, *Irf1*, and *Myd88*). Data are shown as mean, and n are individual animals.

To address this question, we used transcription factor (TF) foot-printing analysis to quantitatively evaluate the occupancy of TFs to the open chromatin regions by identifying smaller regions (10-20 base pairs) of occupied chromatin within the broader open chromatin regions (200-500 base pairs) and identifying the matching TF motif^30,31^. We followed the strategy from established studies that examined how transcriptional gene expression is regulated by TFs through this analysis based on chromatin accessibility data^31–34^. We identified 83 TFs with differential occupancy between MIA and CON microglia with 76% showing less occupancy in MIA compared with CON microglia including Gata4, Smad3, and Stat1 (**Fig. 3d**), which are important TFs that regulate the immune response, particularly the interferon response^35–38^. There is a precedence from previous studies that showed a nonlinear relationship between open chromatin and TF occupancy in other biological systems^39–41^. To confirm the ATAC TF occupancy results indicating impaired interferon signaling, we found that the *Stat1* gene and several Stat1 target genes (*B2m*, *Il1b*, *Irf1*, *Myd88*) consistently showed a reduced gene expression in MIA microglia after LPS treatment (**Fig. 3e**). In summary, while the chromatin is more open in adult MIA microglia (compared with CON microglia) upon LPS treatment, there are fewer TFs occupying these open regions, which is likely to underlie the blunted immune response program at the molecular level.

### Rescue of MIA Blunted Immune Response by Prenatal Microglia Replacement

We next tested whether the long-term microglia impairment may be ameliorated by ablating the MIA-impaired microglia to allow naïve microglia to re-infiltrate the developmental brain after the MIA stressor. To accomplish the prenatal microglia replacement, we treated pregnant dams with a Csf1r antagonist, PLX5526, between E9.5 and E12.5 to effectively eliminate embryonic myeloid cells (**Ext Data Fig. 6**). The short treatment of the Csf1r antagonist did not impact the MIA maternal immune response as measured by serum IL-6 protein expression (**Ext Data Fig. 6f**). In adulthood, we compared the gene expression that changed in response to LPS (LPS response genes) among CON microglia, MIA microglia, repopulated microglia after MIA (repopulated MIA microglia), and repopulated microglia in CON. We found that the LPS response in the repopulated MIA microglia showed an amelioration of the blunted immune response by gene expression (**Fig. 4a-g, Ext Data Fig. 7**). MIA microglia, repopulated MIA microglia, and repopulated CON microglia showed slopes for the fold change linear regression of 0.74, 0.86, and 0.90, respectively, indicating that the LPS response of the repopulated MIA microglia are more similar to the LPS response of CON microglia (0.90) than that of MIA microglia (0.74) (**Fig. 4a-g**).

**Figure 4:**
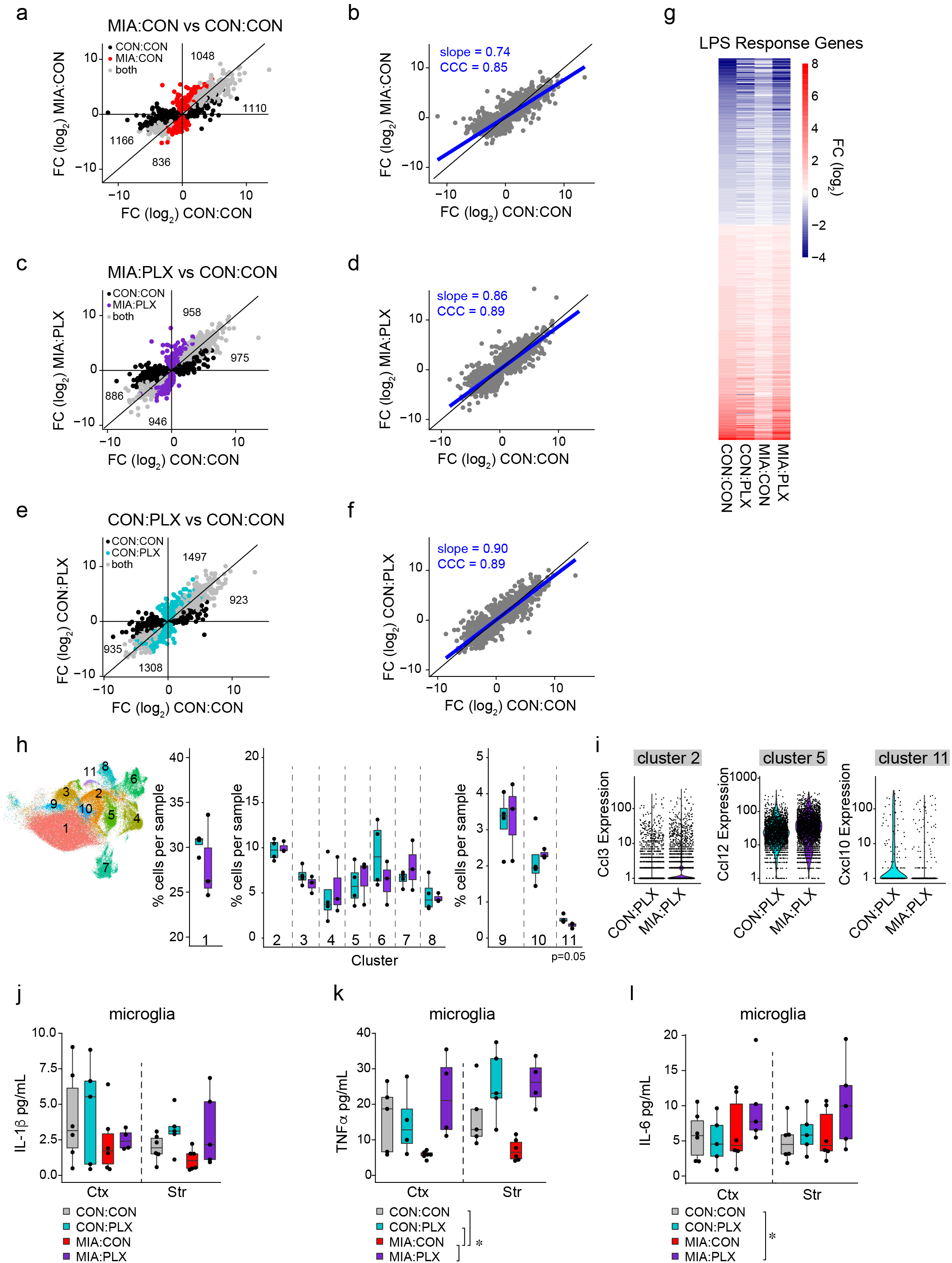
Amelioration of microglia blunting after prenatal replacement. CON:CON, CON mice with control treatment; MIA:CON, MIA mice with control treatment; CON:PLX, CON mice with PLX treatment; and MIA:PLX, MIA mice with PLX treatment. **a-b**, Plot (a) and linear regression (b) of the FC for the LPS-induced DEGs in both CON:CON microglia and MIA:CON microglia (grey), only CON:CON microglia (black), or only MIA:CON microglia (red) from the whole Str. Numbers indicate the total number of LPS response genes in each section. The black line indicates the y=x line and the blue line is the linear fit line (slope = 0.74 and CCC = 0.85). **c-d**, Plot (c) and linear regression (d) of the FC for the LPS-induced DEGs in both CON:CON microglia and MIA:PLX microglia (grey), only CON:CON microglia (black), or only MIA:PLX microglia (purple) from the whole Str. Numbers indicate the total number of LPS response genes in each section. The black line indicates the y=x line and the blue line is the linear fit line (slope = 0.86 and CCC = 0.89). **e-f**, Plot (e) and linear regression (f) of the FC for the LPS-induced DEGs in both CON:CON microglia and CON:PLX microglia (grey), only CON:CON microglia (black), or only CON:PLX microglia (cyan) from the whole Str. Numbers indicate the total number of LPS response genes in each section. The black line indicates the y=x line and the blue line is the linear fit line (slope = 0.90 and CCC = 0.89). **g**, Heatmap of the log_2_ FC for LPS response genes in CON:CON, CON:PLX, MIA:CON, MIA:PLX. **h,** UMAP of ∼120,000 microglia grouped into 11 clusters. Differential abundance analysis quantifying the percent of cells from each sample in each cluster. n are individual animals. Data are shown as median ± interquartile range and analyzed using a t-test. **i,** Gene expression (log_10_) for the distinct marker genes for clusters 2, 5, and 11 in CON and MIA microglia after prenatal replacement. **j-l**, Protein expression of cytokines IL-1β (j), TNFα (k), and IL-6 (l) in microglia from CON:CON (grey), CON:PLX (cyan), MIA:CON (red), and MIA:PLX (purple). n are individual animals. Data are shown as median ± interquartile range and analyzed using two-way ANOVA with emmeans post-hoc test. * p<0.05.

The amelioration of the blunted immune response in repopulated MIA microglia was also addressed at the single cell level (**Fig 4h-i, Ext Data Fig 5**). We found the repopulated MIA microglia did not cluster distinctly from the healthy CON microglia (**Ext Data Fig 5c**). Furthermore, the abundance assay to quantify the proportion of cells from each sample to each cluster found the repopulated MIA microglia showed a rescued contribution to cluster 2 (*Ccl3+*) and cluster 5 (*Ccl12+*) (**Fig 4h-i**). However, cluster 11 (interferon cluster, *Ifit2+ and Cxcl10+*) still showed a slight reduction in the repopulated MIA microglia (**Fig 4h-i**). These data indicate that the re-populated cells are indistinguishable from CON (endogenous) microglia, and the repopulated MIA microglia ameliorated the reduced contribution to the proinflammatory clusters.

The amelioration of the blunted immune response in the repopulated MIA microglia was confirmed at the protein level using IL-1β, TNFα, and IL-6 (**Fig 4j-l**). Adult MIA microglia upon LPS treatment showed reduced protein expression of TNFα, a trend towards decreased lL-1β, but no change in IL-6, whereas the re-populated MIA microglia did not show such deficits (**Fig. 4j-l**). These protein data are consistent with the blunted immune response in the transcriptional data in which the blunting in MIA microglia is rescued by the prenatal microglia replacement.

### Cell Non-Autonomous Contribution of MIA Microglia to Astrocytes and Neurons

To evaluate cell non-autonomous impacts of MIA microglia on the surrounding glia and neurons, we evaluated cytokine protein expression of the astrocytes and non-microglial/non-astrocytic (negative) cells after LPS treatment. In contrast to the reduced protein expression in MIA microglia (**Fig. 4j-l**), astrocytes and non-microglial/non-astrocytic (negative) cells showed increased expression of IL-6 in MIA indicating that astrocytes have an enhanced immune response (**Fig. 5a-b**). IL-1β and TNFα were undetectable in the astrocytes and non-microglial/non-astrocytic (negative) cells after LPS treatment. Prenatal microglia replacement normalized the pathological change of astrocytes (**Fig. 5a-b**), indicating that the impaired microglia reactivity likely underlies the changes in astrocytes of MIA offspring.

**Figure 5:**
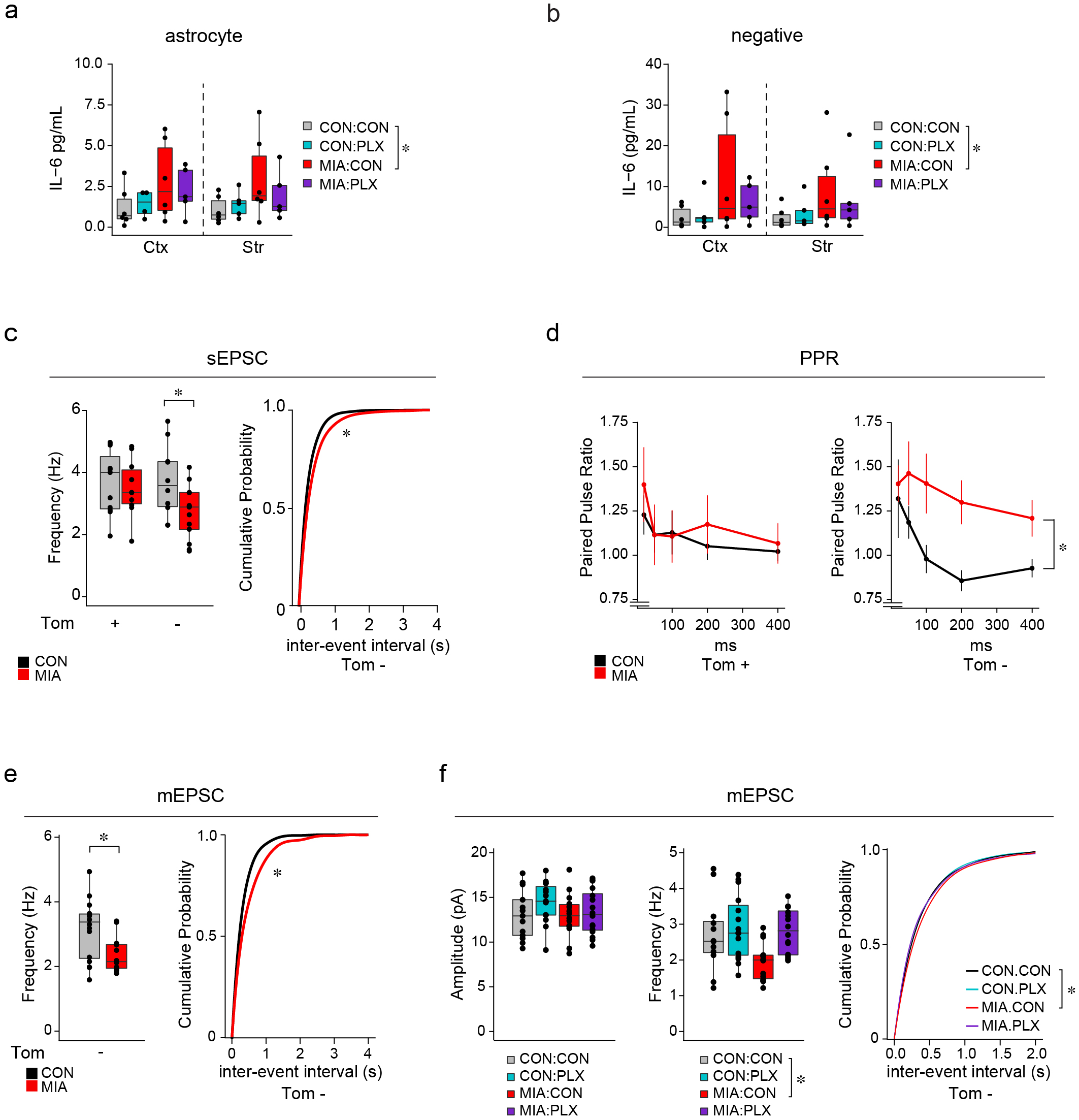
Rescue of astrocyte and neuronal phenotypes in MIA using prenatal microglia replacement. **a-b**, Protein expression of cytokine IL-6 in astrocytes (a) and non-astrocyte/non-microglial cells (negative) (b) from CON:CON (grey), CON:PLX (cyan), MIA:CON (red), and MIA:PLX (purple) offspring. n are individual animals. Data are shown as median ± interquartile range and analyzed using two-way ANOVA with Dunnett’s post-hoc test. * p<0.05. **c**, Spontaneous excitatory postsynaptic current (sEPSC) from Drd1a-TdTomato+ (D1R, Tom+) and Drd1a-TdTomato- (D2R, Tom-) medium spiny neurons (MSNs). n are individual cells. Data are shown as median ± interquartile range. Data was analyzed by t-test with cumulative distribution analyzed by Kolmogorov–Smirnov (KS) test. **d**, Paired pulse ratio (PPR) with a 20-400 msec interstimulus interval from Tom+ (D1R) and Tom- (D2R) MSNs. n are individual cells. Data are shown as mean ± s.e.m. and analyzed by repeated measures two-way ANOVA. **e,** Miniature EPSC (mEPSC) frequency from D2R (Tom-) MSNs. n are individual cells. Data are shown as median ± interquartile range and analyzed by t-test with cumulative distribution analyzed by KS test. **f**, mEPSC amplitude (left) and frequency (center) of D2R MSNs (Tom-) from CON:CON (grey), CON:PLX (cyan), MIA:CON (red), and MIA:PLX (purple) with a cumulative distribution plot for mEPSC frequency (right). n are individual cells. Data are shown as median ± interquartile range and analyzed using a linear mixed effect model and KS test for the cumulative distribution. * p<0.05.

To evaluate the cell non-autonomous impact of MIA microglia on brain development and neuronal function, we evaluated the function of medium spiny neurons (MSNs) in the ventral striatum of adult MIA offspring by electrophysiological recordings. Dopamine dysfunction was previously identified in MIA and the striatum is a major target of the dopamine neuron circuitry^2,42^, but to the best of our knowledge, no study has addressed MSN dysfunction in MIA offspring at the electrophysiological level. We found a decreased frequency of the spontaneous excitatory postsynaptic current (sEPSC) in dopamine receptor type-2 (D2R)-MSNs, but not in D1R-MSNs (**Fig. 5c, Ext Data Fig. 8a**). The sEPSC amplitude was unchanged in both types of MSNs (**Ext. Data Fig. 8b**). We found no change in the spine density in MSNs of the ventral striatum (**Ext. Data Fig. 8c**). However, we found an increase in the paired pulse ratio of D2R-MSN, but not D1R-MSNs (**Fig. 5d**), suggesting a decrease in the presynaptic probability of vesicle release onto D2R-MSNs in adult MIA offspring. To ensure the decreased presynaptic release probability was independent of presynaptic cell excitability, we measured miniature EPSCs (mEPSCs) in D2R-MSNs and found a decrease in the frequency, without any change in the amplitude (**Fig. 5e, Ext Data Fig. 8d**).

Finally, we aimed to determine whether the reduced connectivity of D2R-MSNs was causally related to MIA microglia during the developmental trajectory. To address this question, we performed mEPSC recordings from CON and MIA offspring after prenatal microglia replacement. We found that the reduced mEPSC frequency in MIA offspring was rescued to the control levels after prenatal microglia replacement with no change in amplitude (**Fig. 5f**). In addition, the microglia replacement alone did not alter the physiological properties of the D2R-MSNs in CON offspring (**Fig. 5f**). These results imply that MIA microglia causally underlie the dysfunctional connectivity of the ventral striatal circuit with a decrease in the glutamatergic presynaptic release probability specifically onto D2R-MSNs.

## Discussion

The present study addresses a fundamental biological question of how infiltrating microglia during early brain development are affected by their prenatal environment that may influence neuronal network formation. We demonstrated that prenatal immune stress diminishes microglia reactivity, which results in a functional impairment of activation to subsequent stressors. The microglia dysfunction may be masked at baseline but is revealed when microglia are needed to respond to either intrinsic (neonatal) or extrinsic (LPS) stimuli. We propose a new working model in which prenatal immune stress acts as a tolerizing stimulus leading to a change in the chromatin structure, TF occupancy, and transcriptional regulation of gene expression. Additionally, our data accounts for an enigma in clinical studies in which patients with schizophrenia show increased cytokine levels in cerebrospinal fluid, but brain imaging data suggests no major increase or even a decrease in cellular immune activation^43–47^. We determined a hypoactivation of microglia while showing an increased inflammatory state of astrocytes in adult MIA offspring. Therefore, our data emphasizes the importance of evaluating additional cells, to microglia, as a source of immune molecules in disease.

An outstanding question for future studies is to carefully address the impact of sex on microglia functional deficits since male and female microglia show different immune responsiveness^48–50^ and developmental and psychiatric disorders also show sex differences^51,52^. While we show some cursory evidence that the blunting phenotype is more robust in males (**Ext Data Fig. 9a-d**), this needs to be explored in greater detail in future studies. Another interesting extension is on the specificity of the D2R-MSN physiological change in MIA in the ventral striatum. A recent study showed the critical role of microglia in regulating D1R-MSN circuit development^3^ and we expect similar mechanisms are likely involved in the D2R-MSN circuitry. Finally, further research is required to investigate the microglia states induced by LPS including the functional roles, temporal regulation, and spatial allocation of these activation states.

**Supplementary Information** is available in the online version of the paper.

## Acknowledgements

We thank Travis Faust, Margaret McCarthy and Jonathan Ling for critical reading of the manuscript and Yukiko Lema for assistance with the figures. We thank Hao Zhang for assistance with flow cytometry and cell sorting. We thank Gabrielle Cannon for guidance with single cell library preparation and RNA sequencing. We would like to acknowledge support for the statistical analysis by Leah Jaeger in the National Center for Research Resources and the National Center for Advancing Translational Sciences (NCATS) of the National Institutes of Health through Grant Number 1UL1TR001079. We thank Romain Nardou for advice on electrophysiology. We appreciate the Johns Hopkins Solomon H. Snyder multi-photon imaging core for microscopy and software support (NIH P30 NS050274). We also acknowledge the Plexxikon company for providing the PLX5526 compound. This work was supported by NIH grants (P50MH094268, RO1MH105660, and RO1MH107730), and foundation grants from S-R/RUSK, Brain & Behavior Research Foundation (NARSAD) and the Stanley Center.

## Author Contributions

L.N.H. and A.S. conceived the study. L.N.H performed experiments with help from M.P., M.K., A.C., C.D. L.B. and A.R.. L.N.H. and K.A. performed electrophysiology experiments with help from G.D.. L.N.H., E.V., and L.A.G. performed single cell RNA sequencing experiments and data analysis. L.N.H. and A.S. wrote the manuscript with input from all other authors. A.S. and S.K. supervised the study.

## Author Information

Reprints and permissions information is available at www.nature.com/reprints. The authors declare no competing financial interests. Readers are welcome to comment on the online version of the paper. Correspondence and requests for materials should be addressed to A.S.

## Methods

### Animal husbandry

All animal procedures were approved by the IACUC at Johns Hopkins University. Animals were maintained in individually ventilated cages with a 14-hour light cycle and allowed food and water *ad libitum* in a temperature- and humidity-controlled facility (22 ± 3 °C, 45 ± 5%). Mice were maintained on a standard diet of 18% protein (sterilized 2018SW, Envigo). C57BL/6J mice were purchased from Jackson labs (00664), dopamine receptor D1alpha-TdTomato (Drd1α-TdTomato) mice were kindly shared by Dr. Gul Dölen backcrossed to C57BL/6J mouse line. Male mice were used for breeders and mated with wild-type females to generate CON and MIA pups.

### Generation of MIA and microglia replacement

To generate MIA pups, virgin females (C57BL/6J, 7-12 weeks of age) were plug checked each morning after mating and the appearance of a vaginal plug was demarcated as embryonic day 0.5. The timed-pregnant females were delivered polyinosine-polycytidylic acid (PIC, Sigma, P9582, P0913) by intraperitoneal injection (IP) between 8:00 and 11:00 am on E9.5. PIC or saline was delivered 10-20 mg/kg at a volume five times the animal weight (5 mL/kg). We used E9.5 as our MIA time-point because this is when the microglia infiltrate the neuroepithelium^4^. To ensure efficacy of the maternal immune event, we monitored IL-6 protein expression in the maternal serum levels 3-4 hours post injection. Maternal immune responses below 1000 pg/mL were excluded (**Extended Data Fig. 9e**). Each new batch of PIC was quality controlled to ensure it elicited a comparable immune response, and we found that a serum response greater than 20 ng/mL led to a severe loss of pup viability (**Extended Data Fig. 9e**). For rescue experiments, pregnant dams were placed on PLX5622 (Plexxikon, colony stimulator factor 1 receptor antagonist) or control chow for 3 days between E9.5 to E12.5. On E12.5 mice were returned to house chow.

### LPS injection

Lipopolysaccharide (LPS, Sigma, L4391) was reconstituted in saline to a concentration of 600,000 EU/mL (EU, endotoxin units). Peripheral LPS was injected by IP at a dose of 4500 EU/g body weight. Serum IL-6 was measured at 18 hours post injection to ensure responsiveness.

### Blood draw

Peripheral immune response was monitored to establish a threshold for responsiveness. To collect blood, mice were immobilized and the mandibular/facial vein was pricked with a 4 mm lancet or the tail end snipped to collect blood drops into a 1.5 mL tube. The blood was allowed to clot for at least 15 min then centrifuged at 9,000 RPM for 5 min twice and the supernatant was collected and frozen until assayed.

### ELISA

Cytokines in the serum and cell culture supernatant were measured using the eBioscience Ready-Set-Go ELISA kits according to the manufacture instructions. Briefly, 96-well plates were coated with capture antibodies for either IL-6 or TNFα overnight. The plates were washed in buffer, blocked for 1-2 hours with 1X assay diluent, and then diluted standards and samples were incubated in the plate overnight. Finally, the plate was washed, treated with detection and HRP antibodies, and developed using TMB reaction with a nitric acid stop solution. The colorimetric signal was measured on a plate reader at 450 nm subtracted by 570 nm (Bio-Rad). Maternal serum was diluted 1:100, cell culture supernatants were diluted 1:50, and 18-hour LPS response serum was diluted 1:10.

Cell homogenates were measured using a more sensitive ELISA kit (V-Plex, Mesoscale Discovery). Cells were isolated by sequential MACS sorting for microglia (CD11b) then the flow-through was stained with ASCA2 to isolate astrocytes, and the final flow-through was collected as the negative fraction. The cells were homogenized using RIPA buffer (150 mM NaCl, 1% NP50, 0.1% SDS, 0.5% sodium deoxycholate, 50 mM Tris-HCl, 1x protease inhibitor) with sonication, insoluble protein was pelleted, and soluble protein was quantified using the Micro BCA Protein Assay (Thermo). IL-6, TNFα, and IL-1β were measured according to the manufacture instructions. Briefly, the plate was washed 3 times before adding the diluted samples (1 μg/μl) and standards for 2 hours. The multiplex plate was washed 3 times and the detection antibody was incubated for 2 hours. Finally, the plate was washed 3 times, 2X read buffer applied, read on a MSD plate reader, and analyzed on the MSD Workbench software.

### Immunohistochemistry

Embryos or tissue sections were collected and fixed overnight in 4% paraformaldehyde then cryopreserved in 15% and 30% sucrose. Embryonic tissue was frozen in OCT in a bath of isopentane cooled in liquid nitrogen and then cryosectioned on a Leica cryostat at 12-15 µm sections. Adult tissue sections were sectioned on a Leica vitrabome at 50 um and stored at 4°C. For immunostaining, sections were rinsed in PBS, blocked in 10% normal donkey serum, and then stained in antibodies overnight (rabbit anti-Iba1, 1:500, Wako and CD68 (Biorad 1:100)). The following day the sections were rinsed in PBS three times, stained with secondary antibodies (Alexa 488; Alexa 555), then counterstained with DAPI, and mounted in aqueous mounting media (fluoromount-g). Images were captured on a Keyence microscope at 10x magnification. High resolution images were collected on a Zeiss AxioObserver D1 at 20x and images were tiled and enhanced with Adobe Photoshop. Confocal images were captured on a Zeiss LSM 880 Confocal. Z-stacks were collected at 0.5 um z-intervals for the entire stack at 63x magnification. Low magnification images were taken at 10x with 2um z-stack interval.

### Microglia Morphology and Density Analysis

All microglia density and morphology analyses were performed blinded to condition. Using Imaris (x64 version 9.5.1), we quantified the microglia density in the striatum. Using the spots tool, microglia were automatically identified (settings: 10 um diameter with background subtraction). All cells were verified by eye before final quantification based on image area and volume. 3-4 images per region (dorsal or ventral striatum) per animal were analyzed. Data was analyzed on the per animal density averages per region and statistical analysis using a three-way ANOVA.

Microglia morphology was analyzed by Sholl analysis in Imaris. Specifically, individual microglia were surface rendered based on the Iba1 channel and individual cell “masks” were cropped for Sholl analysis. Next, using the filament tool individual microglia was traced using default settings (no spine detection and cell body sphere region ∼ 10-11 um) and manually modified if necessary. Finally, microglia and lysosomal volume was calculated using the surface tool. Specifically, the microglia (Iba1 signal) from the individually cropped cells were used to calculate the Cd68 surface volume within the Iba1 volume. The Cd68 signal was rendered using settings: local contrast, diameter = 1um, greater than 5 voxels, filtered to shortest distance to surfaces. The volume of each individual microglia, the volume of total lysosomes in microglia, and the percent of microglia volume containing lysosomes was calculated.

### Microglia *in vitro* culture

The brains from an entire E18 litter (both male and female) were pooled. After meningeal removal, the cortex was collected, hippocampi removed, and the subcortical striatum or cortex collected into separate tubes. The tissue was mechanically homogenized using 18, 22, and 25 gauge needles successively. After filtering, the cell suspension was plated into T75 culture flasks (5 million cells per flask) coated with poly-d-lysine (100 µg/mL, final). The media (DMEM/F12, 10% heat-inactivated FBS, 1% penicillin/streptomycin) was changed every 3-5 days and M-CSF was supplemented (Peprotech 315-02, 5 ng/mL) for 2 weeks. Previous studies isolated microglia by shaking the flasks at 200-250 RPM for 4-5 hours and then collecting and plating the floating microglia cells in the supernatant^53,54^. We found we could get comparable experimental results, but more cell yield when we collected all the cells in the flask with trypsin, and isolated microglia through magnetic sorting with CD11b magnetic beads. We found both methods gave similar results therefore both are combined in the data. To perform the magnetic sorting, flasks were briefly treated with trypsin to release the cells, inactivated with complete media, and washed with magnetic sorting buffer (PBS, 2 mM EDTA, 0.5% BSA). The cells were treated with CD11b magnetic beads for 30 min, washed with magnetic sorting buffer, and isolated on a magnetic column after washing. The cells were plated in a 96-well plate at 40,000 cells per well overnight. The cells were treated with LPS (100 ng/mL) and IFNγ (10 ng/mL) for 12 hours and the cell culture supernatant collected, spun at max speed to remove any debris, and frozen until assayed.

The adult animals (8-10 wk) were deeply anesthetized with ether, perfused with PBS, and the brains were isolated. The meninges were removed and then the frontal cortex and striatum were collected into 2 mL tubes. We pooled two mice of the same sex for each culture and used both males and females. The brain pieces were diced with scissors and then homogenized with papain according to the adult neural dissociation kit (Miltenyi). Briefly, the tissue was treated with papain and DNase at 37°C for 30 min with rotation. The homogenates were pipetted gently and filtered with a 70 µm strainer. The cell pellet was resuspended in debris removal solution (300 µL debris removal, 700 µL PBS), overlaid with 300 µL of PBS, centrifuged at 3000xg for 10 min, debris aspirated, washed with PBS, and centrifuged at 1000xg for 10 min. The cell suspension was then plated in T25 flasks coated with PLL (15 µg/mL). The media was changed every 3-4 days and supplemented with M-CSF (10 ng/mL). After 2 week, the microglia were isolated by magnetic sorting and stimulated as described above. For *in vitro* experiments, cytokine measurements were normalized to the cell density as measured by a one hour Alamar Blue assay according to the manufacture instruction (Invitrogen).

### Microglia isolation

Male mice (8-10 wk) were deeply anesthetized with ether, perfused with PBS, and brains extracted. The meninges were removed and then the frontal cortex and striatum were collected into 2 mL tubes with HBSS on ice. The frontal cortex was collected from a coronal slice anterior to 1.7 mm bregma and dorsal of the corpus callosum, and the anterior striatum was excised from this slice by collecting the tissue surrounding the nucleus accumbens. The next coronal slice was cut at -0.5 bregma and the dorsal and ventral striatum was excised from this slice, excluding the olfactory tubercle, and all striatum pieces were pooled together. P4 mice were not perfused but blood was drained by decapitation. The brain pieces were diced with scissors and then homogenized with enzymes according to the adult neural dissociation kit (Miltenyi) as described above. The cells were resuspended in 0.5% BSA in HBSS. For magnetic sorting anti-CD11b magnetic beads were added for 30 min at 4C. The cells were washed with buffer and isolated on the magnetic column before final collection into RNA lysis buffer. For sequential collection, the flow-through negative cells were collected and treated with ASCA2 magnetic beads for 30 min before washing and again collecting on a magnetic column. The collected astrocytes and uncaptured flow-through cells were both collected and lysed in RNA lysis buffer. For FACS sorting, the cells were incubated with Fc block (eBiosciences 16-0161-81) for 5 min then incubated with antibodies for CD45-APC (BD Biosciences 561018) and CD11b-BV421 (BD Biosciences 562605) for 30 min, the cells were washed with buffer 2 times before filtering and collecting on a MoFlo Legacy. Cells were gated on size, singlet, and propidium iodide negative (live cells), before sorting on CD45-APC low, CD11b-BV421 positive.

### Quantification of embryonic myeloid cells by FACS

Pregnant dams were euthanized by cervical dislocation and pups quickly removed. Brains were extracted on ice and meninges removed. Whole brains from individual animals were placed in a tube with papain and DNase (Miltenyi adult neural dissociation kit) and digested for 20-30 min at 37°C. The solution was pipetted through a 70 µm strainer, and spun at 2500 RPM for 2 min to collect the cells. Debris was removed using a debris removal solution (Miltenyi, as above). The cells were incubated with Fc block (eBiosciences 16-0161-81) for 5 min then incubated with antibodies (CD45-APC BD Biosciences 561018, and CD11b-BV421 BD Biosciences 562605) for 30 min at 4°C in FACS buffer (0.5% BSA, 2 mM EDTA, 25 mM HEPES, in 1x HBSS), washed well and strained though a 40 µm strainer. Cells were gated on size (singlet) and propidium iodide negative (live cells), before quantifying on CD45-APC+, CD11b-BV421+ cells using a MoFlow sorter XDP sorter.

### RNA isolation for RT-PCR and bulk RNA sequencing

RNA was collected from freshly isolated, flash-frozen tissue or sorted cells according to the kit instruction (Qiagen, Micro-RNAeasy, PicoPure RNA isolation Kits). Single strand cDNA synthesis was prepared from purified RNA using oligo-dT and random hexamer primer mix with superscript III or IV according to the manufacture instructions (ThermoFisher Scientific). RT-PCR was performed using SYBR green master mix and 1 µM mix of forward and reverse primers. The samples were run on a 7900 HT real-time PCR machine with cycles (95°C for 10 min; 95°C for 15 sec, 60°C for 30sec, 72°C for 30sec at 40-50 cycles, and 72°C for 10 min). The melting curve was evaluated for single peak and a standard curve was performed to ensure linear amplification of each primer set. The following primer sets were used: *Iba1* (Taqman Mm01132452_g1), *Tnf* (forward: CCC TCA CAC TCA GAT CAT CTT CT, reverse: GCT ACG ACG TGG GCT ACA G), *Gapdh* (forward: TGC AGT GGC AAA GTG GAG ATT, reverse: TTG AAT TTG CCG TGA GTG GA), *Gapdh* (Taqman Mm99999915_g1). Relative expression was calculated by subtracting the average endogenous control (*Gapdh*) Ct values from each target gene Ct value to generate a delta Ct value. The fold change was then calculated by setting the CON cortex sample to one and calculating the 2^-delta Ct^ for each sample. Data was log normalized for analysis.

### Electrophysiology

Acute parasagittal brain slices (250 µm) containing the nucleus accumbens (NAc) core were prepared from 9 to 12-week-old male heterozygous Drd1a-tdTomato transgenic mice. Briefly, the mice were anesthetized with isoflurane and the brains were quickly dissected and placed in ice-cold sucrose solution containing 76 mM NaCl, 75 mM sucrose, 25 mM D-(+)-glucose, 25 mM NaHCO_3_, 2.5 mM KCl, 1.25 mM NaH_2_PO_4_, 0.5 mM CaCl_2_, 7 mM MgSO_4_ finally equilibrated to pH 7.3–7.4 with 310 mOsm. Slices were prepared with a vibratome (VT1200S, Leica) in oxygenated (95% O_2_, 5% CO_2_) ice-cold sucrose solution and recovered in warm (32–34 °C) oxygenated sucrose solution for 30 min. The slices were transferred to warm (32–34 °C) oxygenated artificial CSF (aCSF) containing 125 mM NaCl, 20 mM D-(+)-glucose, 26 mM NaHCO_3_, 2.5 mM KCl, 1.25 mM NaH_2_PO_4_, 1 mM MgSO_4_, 2 mM CaCl_2_ supplemented with 0.4 mM ascorbic acid, 2 mM pyruvic acid, and 4 mM L-lactic acid, finally equilibrated to pH 7.3–7.4 with 315 mOsm, and then cooled to room temperature until recording. Slices were transferred to a submersion chamber and continuously superfused (3–4 mL/min) with warm oxygenated aCSF (32°C) containing 100 µM of picrotoxin while recording. Medium spiny neurons (MSNs) located in anterior-dorsal part of NAc were visually identified with an upright microscope (Olympus BX51WI equipped with 40x objective, 0.8 NA) using either transmitted light with IR-DIC optics or epifluorescence signal through digital CMOS camera (ORCA-Flash 4.0 LT, Hamamatsu). For the whole-cell patch, microglass recording electrodes (3–5 MΩ) were filled with internal solution containing 120 mM CsMeSO_4_, 15 mM CsCl, 8 mM NaCl, 10 mM HEPES, 0.2 mM EGTA, 10 mM TEA-Cl, 4 mM MgATP, 0.3 mM NaGTP, 0.1 mM spermine and 5 mM QX-314, pH 7.3 with 290 mOsm and recordings were obtained with Multiclamp 700B amplifier (Molecular Devices) controlled by pClampex 10.3 (Molecular Devices), filtered 2 kHz and digitized at 10 kHz with Digidata 1440A (Molecular Devices). Excitatory afferent fibers located between the NAc core and cortex dorsal to the anterior commissure were stimulated with bipolar concentric electrode (FHC) with an intensity which can induce 100–300 pA of EPSCs at -70 mV. Access resistance was monitored throughout the experiments by applying mild hyperpolarizing pulse in each sweep and only the results from the cells changing less than 20% were used for further analysis. I/V relationships were acquired with afferent fiber stimulation while changing holding potential. Rectification indexes for AMPA receptors were calculated by the ratio of the peak amplitudes at +40 mV and -70 mV respectfully. AMPAR/NMDAR ratios were obtained by dividing the peak amplitude of EPSCs at -80 mV (AMPAR EPSCs) with the magnitude of EPSCs at +40 mV at 50 msec after stimulation (NMDAR EPSCs). Paired-pulse ratios were acquired by dividing EPSC2 with EPSC1 obtained from a paired afferent stimulation with the same stimulating intensity by designated inter-stimulus-interval. Spontaneous EPSCs were recorded at -70 mV for 1 min while monitoring access resistance in every 15 sec. Electrophysiology data were displayed off-line with Clampfit software (Molecular Devices). Analysis of spontaneous EPSCs was performed with MiniAnalysis software (Synaptosoft) by automatically detecting negative-going inward EPSCs at a threshold of 5 pA (3x (RMS of noise)^1/2^) and each event was manually inspected. Instantaneous amplitudes and frequencies were acquired to obtain mean value and cumulative graph.

### Spine Counting

Spine counting was performed on golgi stained tissue (FD Rapid GolgiStain Kit). Briefly, male animals were decapitated and brains rapidly extracted. The whole brain was rinsed in water and placed in impregnation solution for 9 days, then transferred to solution C. After 3 days in solution C, the brains were frozen in isopentane, 150 µm sections were collected on a cryostat (Leica) and mounted onto gelatin coated microscope slides. The sections were then rinsed, stained for golgi, and dehydrated before mounting in permount. The spines were quantified with a Zeiss AxioImager microscope and the Neurolucida Software (Microbrightfield). The dorsal and ventral striatum was outlined at low magnification, then cells were selected at 25x, spines were quantified at 100x. We quantified spines that were at 20-200 µm away from the cell body on second to fifth order dendrites. We counted 29-41 cells per group/region, and averaged the spine density per animal. n represents the number of animals.

### Bulk RNA sequencing

Bulk cell RNA sequencing libraries were prepared using a modified Smart-Seq2 protocol^55^. 5 ng of purified RNA were mixed with 2.5 mM dNTP mix, 2.5 µM oligo-dT_30_VN, and 1 U/µL RNasin and incubated at 72°C for 3 min. Then 5.7 µL of the single-cell reverse transctiption mix (1x superscript II buffer, 1 M betaine, 5 mM DTT, 100 U superscript II, 10 U RNasin, 1 µM TSO oligo, 6 mM MgCl_2_) was added to each sample for reverse transcription. The cDNA was amplified with the KAPA HF Ready mix with 9 PCR cycles. The cDNA was cleaned up with AMPure XP beads (Beckman Coulter) at a ratio of 0.7:1, washed with 80% ethanol, and resuspended in 17.5 µL water. The libraries were tagmented for sequencing using the Illumina Nextera XT DNA sample preparation kit. 200-250 pg of cDNA was added to 2.5 µL tagment DNA buffer (Illumina) and 1.25 µL of amplicon tagment mix (Illumina) and incubated at 55°C for 5 min to tagment cDNA. The reaction was neutralized with 1.25 µL of neutralize tagment buffer and incubated at room temperature for 5 min. Then the tagmented DNA was indexed and amplified with 3.75 µL of PCR master mix (Illumina) and 1.25 µL each of i5 and i7 indexing primers (Illumina, diluted 1:5). The samples were indexed with the following PCR cycles: 1 cycle of 72°C for 3 min and 95°C for 30 sec; 12 cycles of 95°C for 10 sec, 55°C for 30 sec, and 72°C for 30sec, and 1 cycle of 72°C for 5 min. The final libraries were purified using AMPure XP beads (Beckman Coulter) at a 0.6:1 ratio, washed with 80% ethanol and resuspended in 12 µL water. The indexed libraries were pooled together at an equal mass ratio for final sequencing on a NovaSeq6000.

### Single cell RNA sequencing

Male mice (8-10 wk) were deeply anesthetized, perfused with HBSS, brains extracted, and frontal cortex and striatum collected into glass tissue homogenizers on ice with homogenization buffer (15 mM HEPES, 5% Trehalose, 500 U/mL DNaseI, 80 U/mL RNasin, in 1x HBSS)^56^. The brain pieces were homogenized 6-7 times, filtered through a 70 µm strainer, and spun at 2500 RPM for 2 min to collect the cells. Debris and myelin was removed using a debris removal solution (Miltenyi, see above). The cells were incubated with Fc block (eBiosciences 16-0161-81) for 5 min then incubated with antibodies (CD45-APC BD Biosciences 559864, and CD11b-BV421 BD Biosciences 562605) for 30 min at 4°C in incubation buffer (0.5% BSA, 2 mM EDTA, 25 mM HEPES, 5% Trehalose, 80 U/mL RNAsin in 1x HBSS), washed well with FACS buffer (0.5% BSA, 2 mM EDTA, 25 mM HEPES, in 1x HBSS), and strained though a 40 µm strainer. Cells were gated on size, singlet, and propidium iodide negative (live cells), before sorting on CD45-APC low, CD11b-BV421 positive using a MoFlo Legacy sorter. Single cells were captured on a 10X Genomics Chromium Platform at a capture target of 6,000 – 10,000 cells. The libraries were pooled and sequenced on a Novaseq6000.

### ATAC Sequencing

Single cells were prepared and FACS sorted as described above. 12,500 - 25,000 cells were collected for each ATAC sample. The ATAC libraries were prepared according to Corces et al. 2017^57^. Briefly, cells were pelleted by spinning at 500g for 15 min at 4C and the supernatant carefully removed. 50 ul of cell lysis buffer was added (10 mM Tris-HCl, 10mM NaCl, 3 mM MgCl_2_, 0.1% NP-40, 0.1% Tween-20, 0.01% Digitonin), pipetted to mix 3-5 times, incubated on ice for 3 min, and washed (10 mM Tris-HCl, 10mM NaCl, 3 mM MgCl_2_, 0.1% Tween-20). The nuclei were spun to pellet at 500 g for 30 min at 4C and the supernatant carefully removed. Then the Tn5 buffer (1X TD Buffer, 1x PBS, 0.1% Tween-20, 0.01 Digotinin, 1 ul Tn5) was added and incubated at 37°C for 1 hour with 1000 RPM shaking. The DNA was collected using the Qiagen Minelute cleanup kit. The DNA was eluted with 11 ul EB Buffer. For library preparation, 10 ul of prepared DNA was mixed with 1.25 uM indexing primers and 1x NEBNext High-Fidelity PCR Master Mix and amplified (72 C 5 min, 98C 30 sec, 5 cycles: 98 10 sec, 63 30 sec, 72 1 min). 5 ul was used for qPCR to test amplification and calculate additional cycles. 6-8 more cycles were run before cleanup with Ampure XP beads. The libraries with quality control checked using a Bioanalyzer then pooled at equal mass for sequencing on a Novaseq6000.

### RNA sequencing analysis

The FASTQ files were first quality checked using FASTQC. The libraries were aligned, assembled, and quantified using RSEM with the STAR aligner (rsem-calculate-expression) using the GRCm38 primary assembly and gencode vM25 primary assembly annotation. Samples were excluded if determined an outlier due to isolated clustering using 2 methods (principal component analysis (PCA) and hclust). Further analysis was performed in R using pipelines for DESeq2^58,59^ and additional modeling with edgeR^60,61^. Gene set enrichment analyses (Broad, GSEA 4.1.0 build 27)^62^ were used for pathway analysis with graphical rendering using cytoscape (v3.8.2)^63^. Gene enrichment for disease association was performed with data from the NHGRI GWAS catalog (v1.0.2, https://www.ebi.ac.uk/gwas/) and included genes with a -log10 p-value >8 and an odds ratio >1 for associations with autism, attention deficit hyperactivity disorder, bipolar disorder, depression, and schizophrenia were counted with the DEGs and non-DEGs using a Chi-square test between the MIA microglia DEGs and the non-DEGs with Bonferroni and Benjamini-Hochberg correction for multiple comparisons.

### Single Cell RNA Sequencing analysis

Single cell libraries were aligned and quantified using the 10X genomics cloud Cell Ranger Count v6.0.0. The libraries were analyzed using Seurat v4^64^. Doublets were estimated with scDblFinder^65^. Low quality cells were filtered using RNA count < 600, feature detected < 500, and percent mitochondrial reads > 10%. The cells from each sample were integrated using SCTransform with glmGamPoi and reciprocal PCA using Seurat^66,67^, dimension reduction with PCA and UMAP, and neighborhood and cluster detection with a resolution of 0.2^68^. Cluster enriched genes were identified using Seurat *FindMarkers*. Differential abundance was analyzed using a t-test and one-way ANOVA for the percent contribution of each sample to each cluster. Samples with low serum IL-6 protein expression were excluded from the differential abundance analysis.

### ATAC sequencing analysis

Library quality was assessed with FASTQC. Libraires were trimmed using cutaddapt^69^ and aligned using bowtie2^70^. PCR duplicates were identified and removed with PICARD MarkDuplicates (Picard Toolkit.” 2019. Broad Institute, GitHub Repository. http://broadinstitute.github.io/picard/). Peaks were called using MACS2 (-f BAMPE -- min-length 100 --max-gap 50)^71^. Coverage and transcriptional start site distances were calculated with DeepTools^72^, and peak counts calculated with FeatureCounts^73^ from subread. Differentially accessible regions were identified using DiffBind^74^ and a permutation test for multiple comparison correction. For footprint analysis, the bam files were shifted for Tn5 correction using alignmentSieve from DeepTools (--ATACshift) and PIQ^33^ for footprint detection using the HOmo sapiens COmprehensive MOdel COllection (HOCOMOCO) (v11) motif database^75^. Transcription factor occupancy thresholds were 0.9 purity score, > 250 total occupancy sites, > 0.5 log_2_ fold change between MIA and CON.

### Statistics

Statistical analyses were performed using the statistical computing program R. Analyses that required subjective measurement were blinded spine density, microglia morphology, and lysosomal content. Exact sample sizes and statistical results are listed in the figure legends and supplemental statistics data table. Data was tested for normality using Shapiro-Wilks test. Normally distributed data was analyzed with a two-tailed t-test and non-normally distributed data was analyzed with a Wilcoxon rank sum test. One- and two-way ANOVAs were performed with pairwise two-sided t-tests with pooled error and Benjamini-Hochberg correction for multiple comparisons. In addition, we used mixed effect linear regression with emmeans to estimate marginal means and include litter as a random effect variable.

### Data Availability

The datasets generated and analyzed during the current study are available from the corresponding author on reasonable request.

## Abbreviations

MIA: Maternal Immune Activation
PIC: polyinosinic-polycytidylic acid
LPS: Lipopolysaccharide
TCA: tricarboxylic acid cycle
ATP: adenosine triphosphate
IL-6: Interleukin 6
TNFα: tumor necrosis factor alpha
E18: embryonic day 18
P4: postnatal day 4
FDR: false discovery rate
MSNs: Medium spiny neurons
D1R: dopamine receptor 1-type MSN
D2R: dopamine receptor 2-type MSN
GSEA: Gene set enrichment analysis
NAc: nucleus accumbens
aCSF: artificial cerebrospinal fluid
IP: intraperitoneal injection
RPM: revolutions per minute
ELISA: enzyme linked immunoassay
HRP: horseradish peroxidase
TMB: 3,3’,5,5’-Tetramethylbenzidine
FBS: fetal bovine serum
M-CSF: Macrophage colony stimulating factor
Cd11b: cluster of differentiation 11b
CD45: cluster of differentiation 45
IFNγ: interferon gamma
PBS: phosphate buffered saline
PDL: poly-d-lysine
PLL: poly-l-lysine
HBSS: Hank’s Balanced Salt Solution
sEPSC: spontaneous excitatory postsynaptic current
mEPSC: mini excitatory postsynaptic current
s.e.m.: standard error of the mean
ns: not significant

---

**Extended Data Figure 1:**
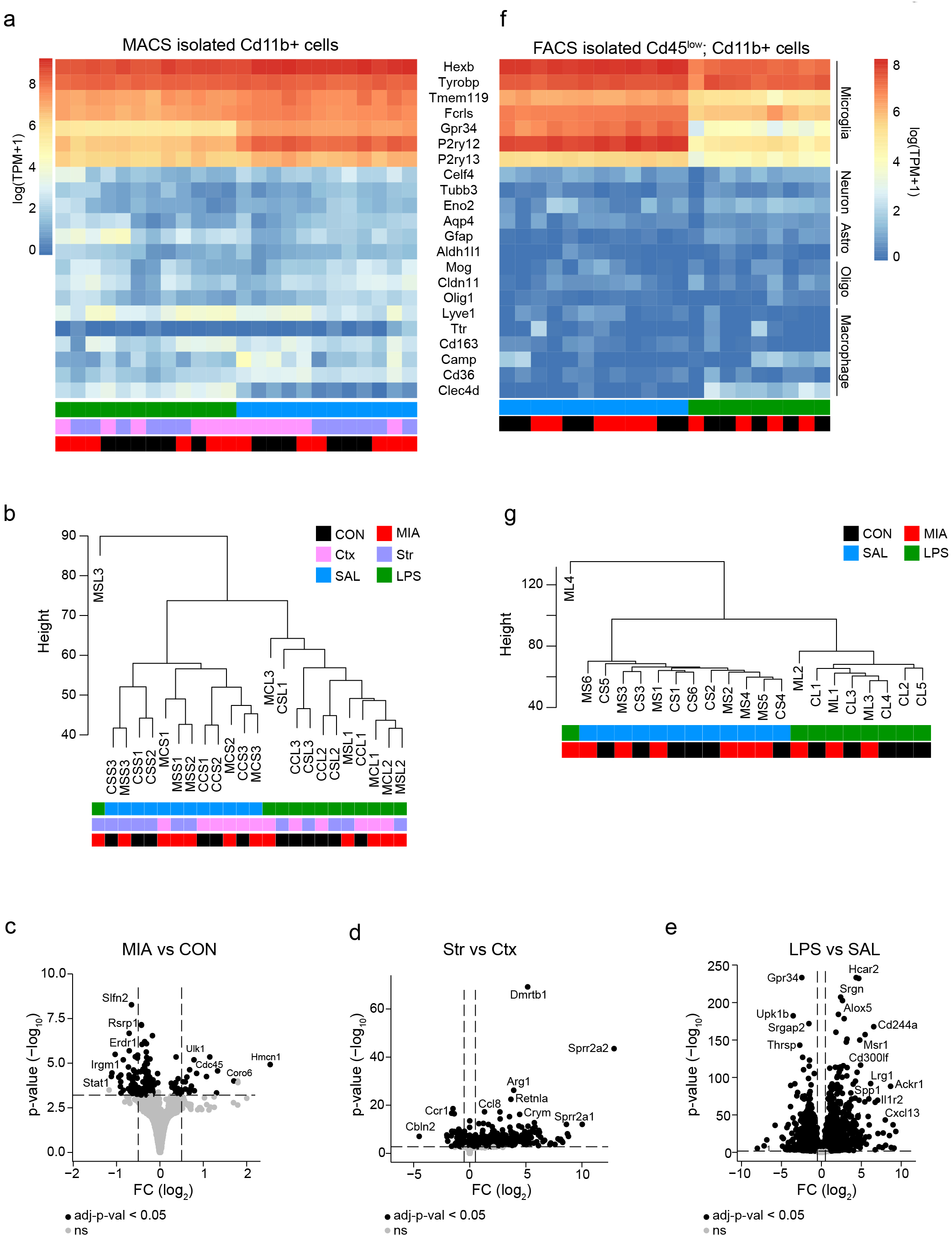
Quality control data of adult bulk microglia RNA sequencing for Figure 1. **a,** Heatmap for the gene expression of cell type specific markers for microglia, neurons, astrocytes, oligodendrocytes, and macrophages from MACS-isolated CD11b+ cells. **b,** Clustering of bulk microglia RNA sequencing samples from (a). One outlier identified (MSL3) and removed from subsequent analysis. Sample naming is as follows: first position M (MIA, red) or C (CON, black), second position C [cortex (Ctx), pink] or S [whole striatum (Str), purple], third position S (SAL, blue) or L (LPS, green), fourth position is animal number. **c-e,** Volcano plots for the DEGs between the main effect of group (MIA versus CON), region (frontal Ctx versus whole Str), and treatment (LPS versus SAL) correcting for the other covariate variables from MACS-isolated CD11b+ cells. Black, q <0.05. **f,** Heatmap for the gene expression of cell type specific markers for microglia, neurons, astrocytes, oligodendrocytes, and macrophages from FACS-isolated CD45^low^;CD11b+ cells from the whole Str. **g,** Clustering of bulk microglia RNA sequencing samples from (f). MS6 was removed from subsequent analysis due to low read count. Sample naming is as follows: first position M (MIA, red) or C (CON, black), second position S (saline, blue) or L (LPS, green), and third position is animal number.

**Extended Data Figure 2:**
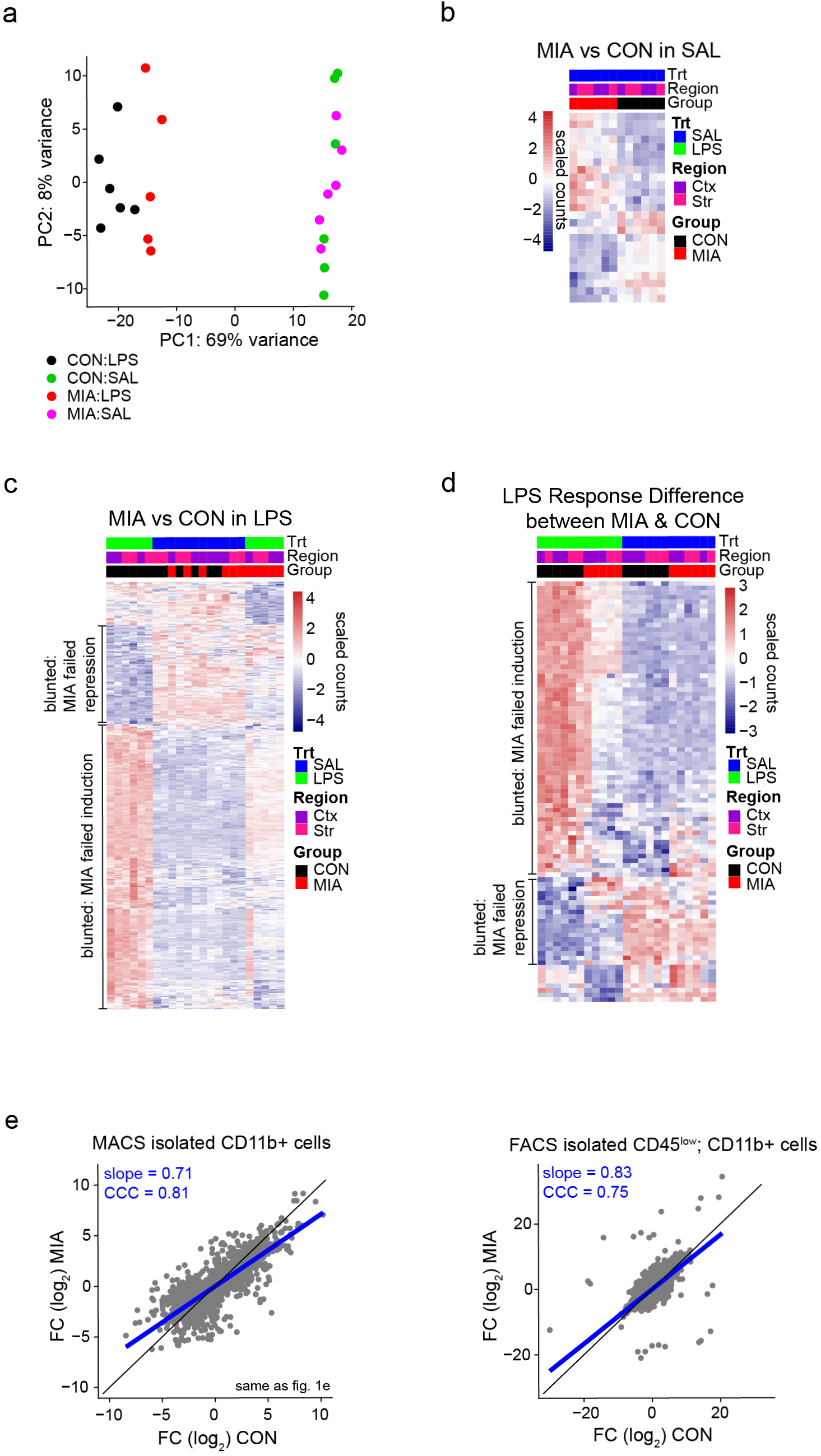
PCA and Heatmaps for DEGs represented in Figure 1. **a,** Principal component analysis (PCA) of CON and MIA microglia after SAL or LPS treatment. **b**, Top 25 DEGs between MIA and CON microglia after SAL treatment. **c**, Top DEGs between MIA and CON microglia after LPS treatment. Blunted LPS-induced genes and blunted LPS-repressed genes are indicated. **d**, Top DEGs between MIA and CON microglia that show an interaction between MIA and LPS (group x treatment). Blunted LPS-induced and LPS-repressed genes are indicated. **e,** Linear regression of the FC for the LPS-induced DEGs from Figure 1e of MACS-isolated microglia (left) in comparison with the FACS-isolated microglia (right). The black line indicates the y=x line and the blue line is the linear fit line with slope equal to 0.71 and 0.83, respectively and CCC equal to 0.81 and 0.75, respectively.

**Extended Data Figure 3:**
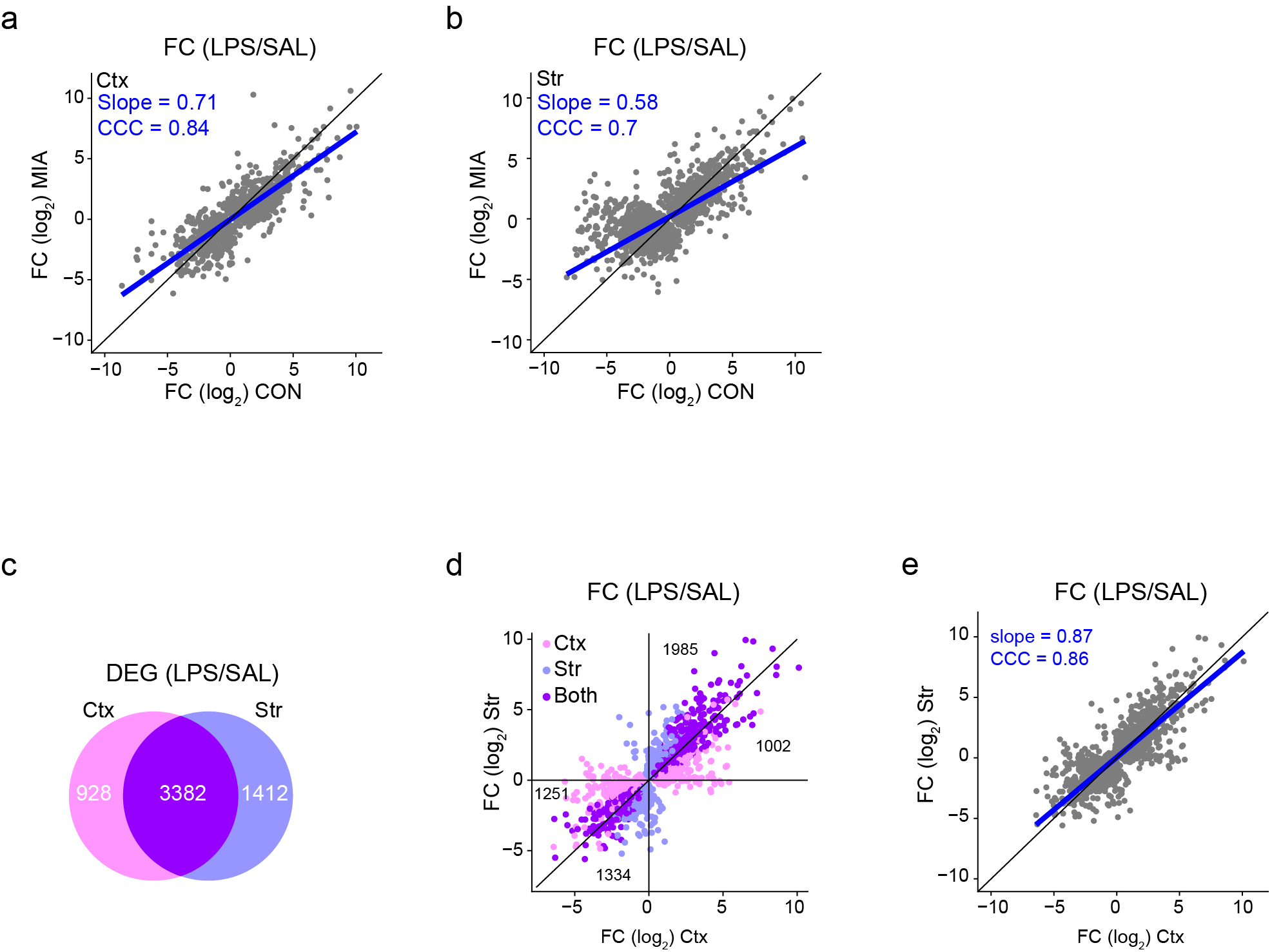
Regional specificity of the diminished microglia immune response after MIA. **a-b**, Regional specificity of the MIA blunted LPS response. Linear regression of the FC for the LPS-induced DEGs between microglia from the frontal Ctx (a) or whole Str (b) with slope equal to 0.71 and 0.58, respectively and CCC equal to 0.84 and 0.70, respectively. **c-e**, Control analysis of the LPS responsiveness between microglia from the Ctx compared with microglia from the Str (these are the mixture of both CON and MIA microglia). **c**, Venn diagram of DEGs (q <0.05) between SAL and LPS in the Ctx microglia (pink) compared with the Str microglia (blue). **d-e,** Plot (d) and linear regression (e) of the FC for the LPS-induced DEGs in both Ctx and Str microglia (purple), only Ctx microglia (pink) or only Str microglia (blue). Numbers indicate the total number of LPS response genes in each section. The black line indicates the y=x line and the blue line is the linear fit line (slope = 0.87 and CCC = 0.86).

**Extended Data Figure 4:**
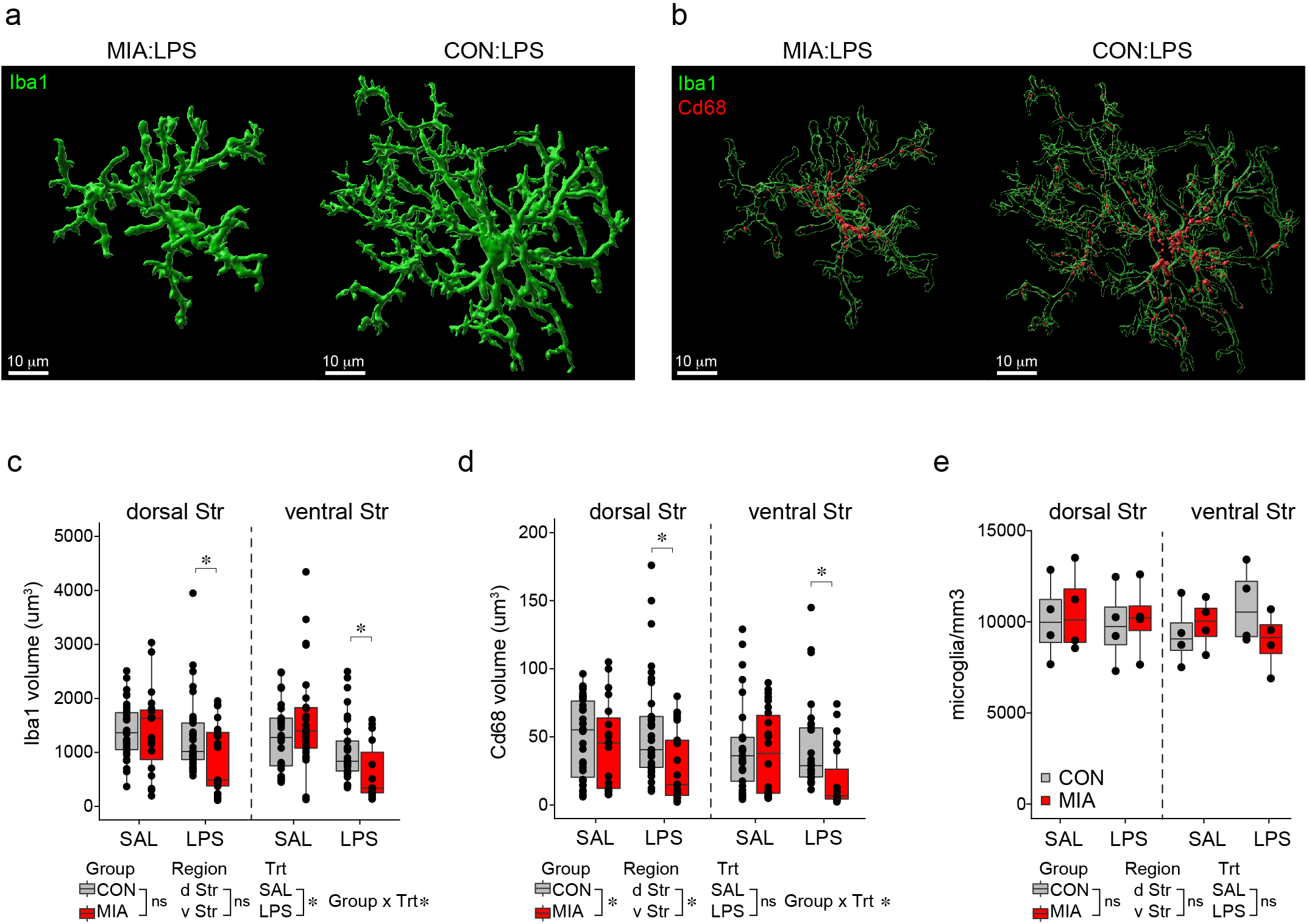
Microglia density and morphology in the striatum after MIA. **a-b**, Representative confocal reconstructions of microglia morphology (a) and CD68 lysosomal content (b) in CON and MIA offspring after LPS treatment. Scale bar = 10 μm **c,** Volume of Iba1+ (microglia) in the dorsal and ventral Str of CON (grey) and MIA (red) microglia after SAL and LPS treatment. n are individual cells. Data are shown as median ± interquartile range and analyzed by a three-way ANOVA with emmeans post-hoc test. * p<0.05. **d**, Volume of CD68 (lysosomes) inside microglia from the dorsal and ventral Str of CON (grey) and MIA (red) offspring after SAL and LPS treatment. n are individual cells. Data are shown as median ± interquartile range and analyzed by a three-way ANOVA with emmeans post-hoc test. * p<0.05. **e,** Microglia density in the dorsal and ventral Str of CON (grey) and MIA (red) offspring after SAL or LPS treatment. n are individual animals. Data are shown as median ± inter quartile range and analyzed by a three-way ANOVA with no significant difference between group, region, or treatment.

**Extended Data Figure 5:**
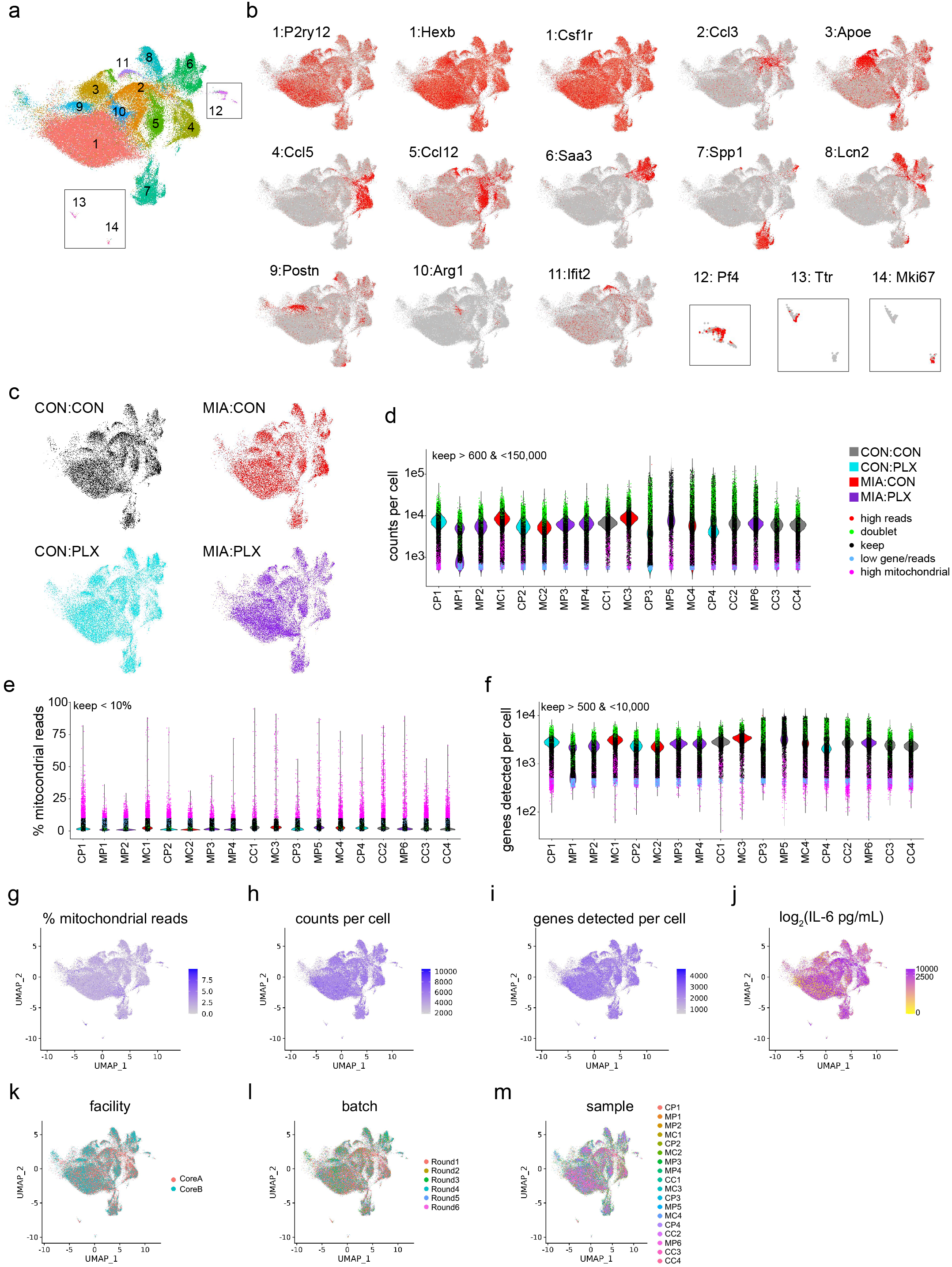
Single cell RNA sequencing of LPS-activated striatal microglia from CON and MIA offspring with prenatal microglia replacement. **a**, UMAP of ∼120,000 microglia grouped into 14 clusters. Note: 3 clusters were removed due to markers for poor cell quality. **b**, Markers enriched for each cluster. Cluster 1 is homeostatic microglia expressing *P2ry12*, *Hexb*, and *Csf1r*. Cluster 2 expresses *Ccl3*, cluster 3 expresses *Apoe*, cluster 4 expresses *Ccl5*, cluster 5 expresses *Ccl12*, cluster 6 expresses *Saa3*, cluster 7 expresses *Spp1*, cluster 8 expresses *Lcn2*, cluster 9 expresses *Postn*, cluster 10 expresses *Arg1*, cluster 11 expresses *Ifit2* and *Cxcl10*. Cluster 12 and 13 are macrophage clusters expressing *Pf4* and *Ttr*, respectively. Cluster 14 is a proliferative cluster expressing *Mki67* and *Top2a*. **c**, Clustering distribution of microglia from CON (CON:CON, black), MIA (MIA:CON, red), prenatal microglia replacement (CON:PLX, cyan), and MIA with prenatal microglia replacement (MIA:PLX, purple). **d-f**, Filtering of cells (∼147,0000) for low counts per cell (d, blue dots < 600 counts), low number of total genes detected per cell (f, blue dots, < 500 genes), high mitochondrial content (e, magenta, > 10% mitochondrial), doublet detection (green dots) or high read count (red dot, > 150,000). **d**, RNA counts per cell for each sample; cells were retained with counts > 600 and < 150,000. **e**, Percent mitochondrial reads per cell for each sample; cells were retained with < 10% mitochondrial content. **f**, Number of genes detected per cell for each sample; cells were retained with > 500 genes and < 10,000 genes detected. Cell doublets were also removed and indicated by green dots in d-f. **g-m**, UMAP of cells after filtering. **g**, UMAP of percent mitochondrial reads per cell. **h**, UMAP of RNA counts per cell. **i**, UMAP of detected genes per cell. **j**, UMAP of serum IL-6 expression quantified in each sample. **k**, UMAP of batch effect for processing facility. **l**, UMAP of batch effect depending on date of processing. **m**, UMAP of individual samples processed. n are individual cells.

**Extended Data Figure 6:**
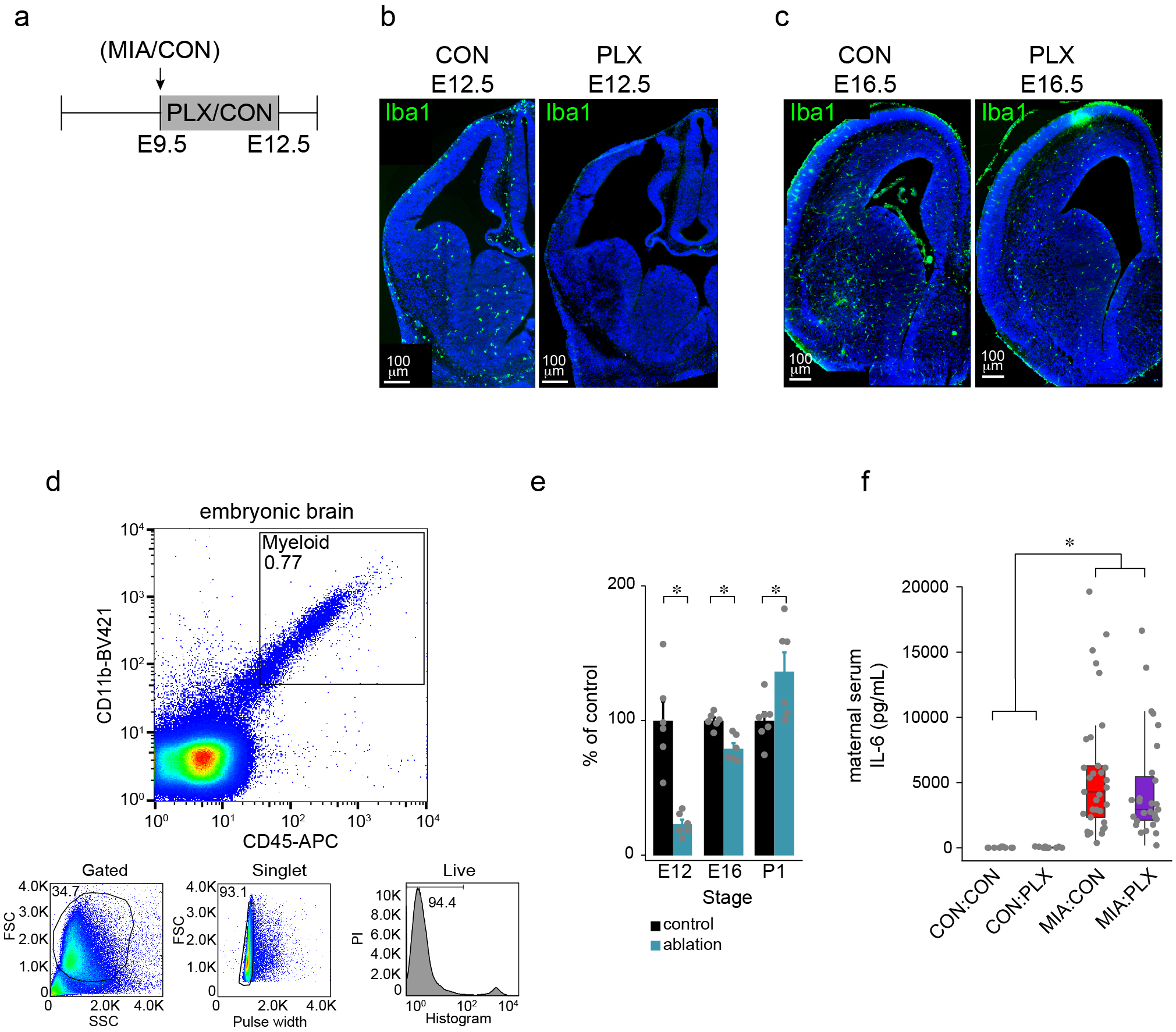
Ablation and repopulation efficiency of embryonic microglia. **a,** Timeline of PLX5622 (PLX) treatment. **b-c,** Representative images of E12.5 (b) and E16.5 (c) brain slices with high efficiency of ablation after 3 days of ablation treatment (b) and 4 days of repopulation (c). Green= Iba1, Blue=Dapi, scale bar = 100 μm. Images are representative from 4-9 individual animals per condition. **d,** FACS sorting plot for quantification of repopulation of brain myeloid cells after gating for live singlets, the CD11b+CD45+ cells were quantified relative to control animals. FSC=forward scatter, SSC=side scatter, PI=propidium iodide. **e,** Quantification of microglia ablation and re-infiltration. n are individual animals. Data are shown as mean ± s.e.m and analyzed using a t-test. *p<0.05. **f**, IL-6 protein expression in maternal serum 3 hours after polyinosinic-polycytidylic acid (PIC) or saline injection with and without prenatal PLX treatment. n are individual animals. Data are shown as median ± interquartile range and analyzed using a two-way ANOVA with Tukey post-hoc test. * p<0.05.

**Extended Data Figure 7:**
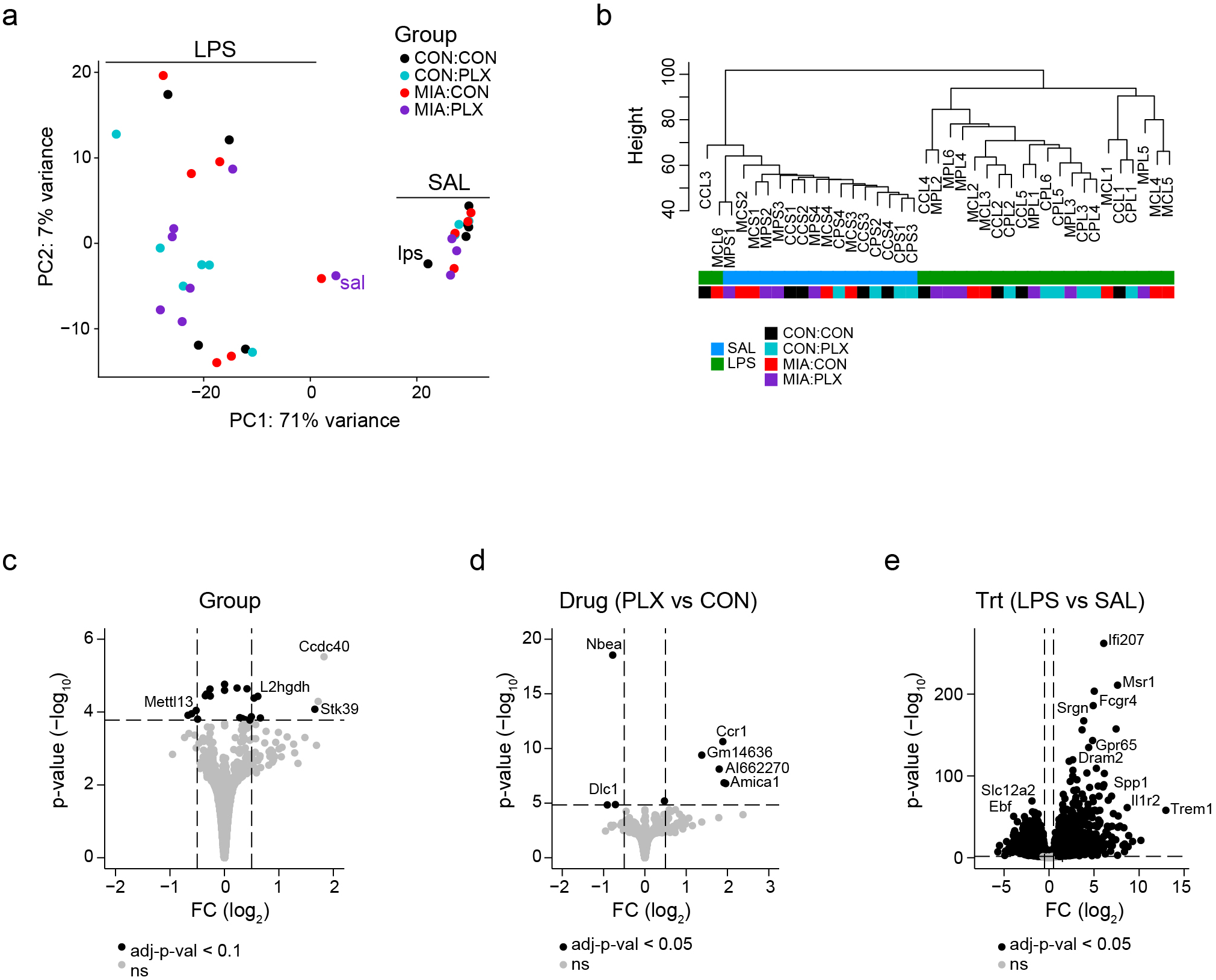
Rescue of microglia blunting in MIA offspring using prenatal microglia replacement. **a,** PCA of microglia from CON:CON (black), CON:PLX (cyan), MIA:CON (red), and MIA:PLX (purple) from the whole Str. Two samples were excluded due to incorrect clustering, each labeled accordingly. **b**, Clustering of bulk microglia RNA sequencing samples from (a). **c-e**, Volcano plots for the DEGs between the main effect of group (c, among CON:CON, CON:PLX, MIA:CON, and MIA:PLX), PLX treatment (d, PLX versus CON), and LPS treatment (e, SAL versus LPS) correcting for the other covariate variables. Black q <0.05.

**Extended Data Figure 8:**
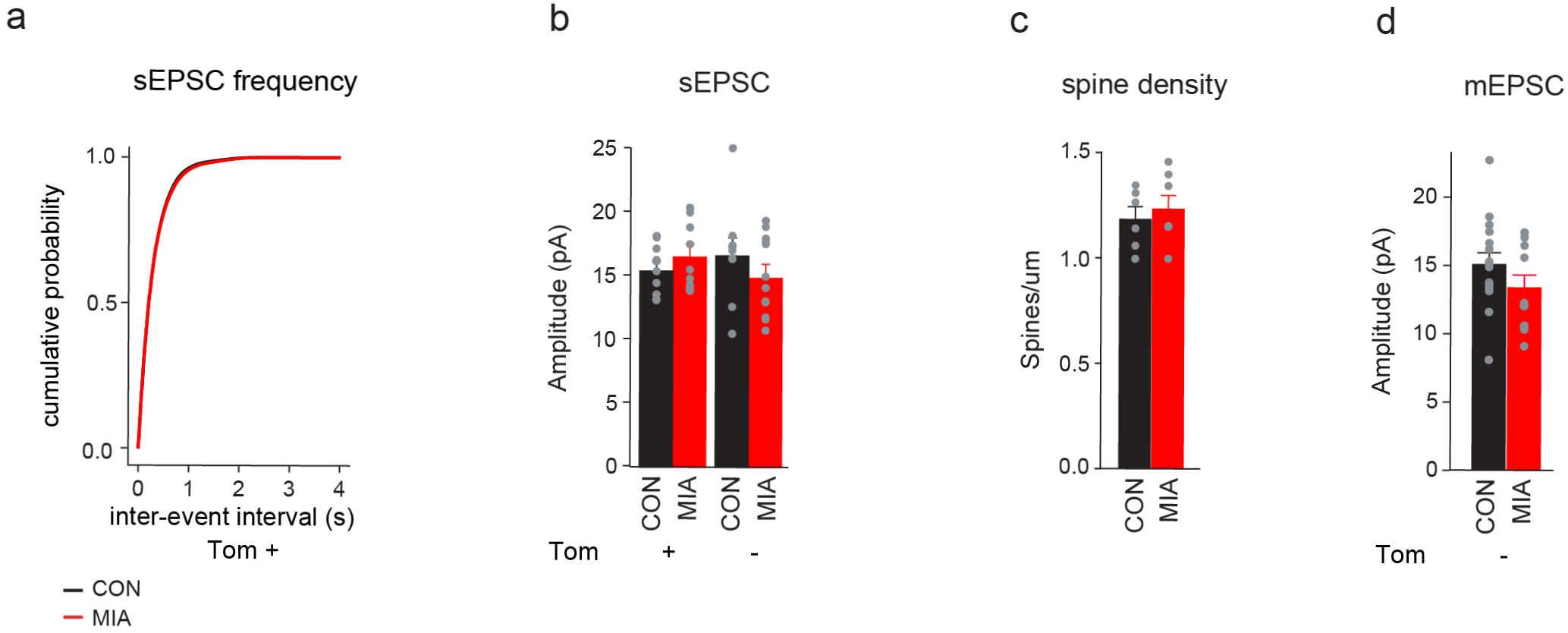
Rescue of neuronal electrophysiological phenotypes in MIA offspring using prenatal microglia replacement related to Figure 5. **a,** Two-sample KS test for the cumulative distribution of the inter-event intervals of the sEPSC frequency from Tom+ (D1R) in Fig. 5c. **b,** sEPSC amplitude of Tom+ (D1R) and Tom- (D2R) medium spiny neurons (MSNs) in Fig. 5c. n are individual cells. Data are shown as mean ± s.e.m and analyzed using t-tests. **c,** Spine density of striatal MSNs in the ventral Str. n are individual animals. Data are shown as mean ± s.e.m and analyzed using a t-test. **d**, mESPC amplitudes of Tom- (D2R) MSNs analyzed in Fig. 5e. n are individual cells. Data are shown as mean ± s.e.m and analyzed using a t-test.

**Extended Data Figure 9:**
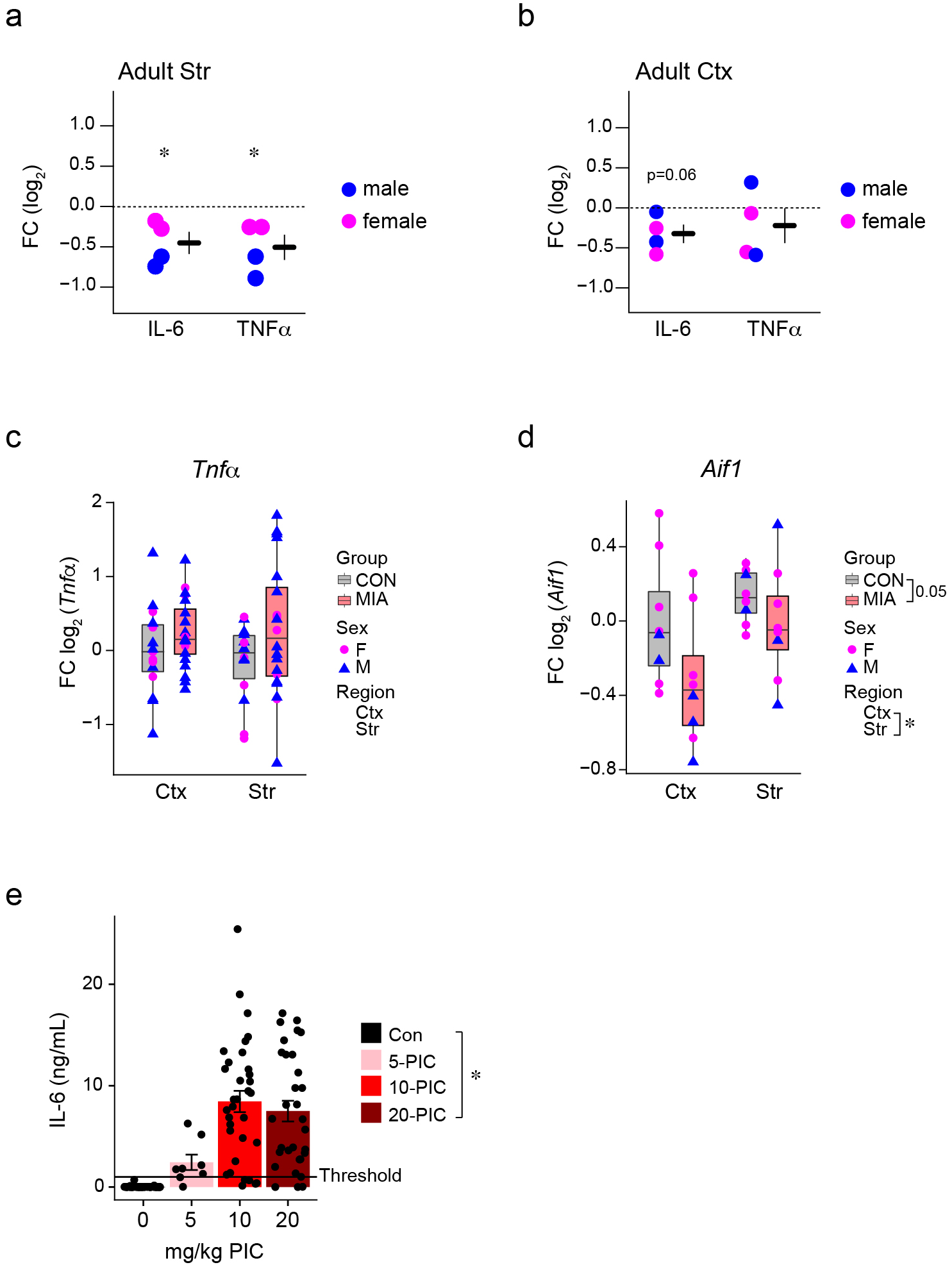
Sex effect in MIA and three-hour dose response in maternal serum after MIA. **a-b**, Quantification of secreted IL-6 and TNFα from LPS-stimulated microglia *in vitro* at adulthood from the Str (a) and frontal Ctx (b) from Fig. 2a-b indicating the sex [male (blue) and female (pink)] of each offspring. n are individual culture experiments. Data are shown as mean ± s.e.m. with each experiment normalized to CON and analyzed using a one-sample t-test. *p<0.05. **c-d**, Baseline gene expression analyzed by qPCR for *Tnfa* (c) and *Aif1* (d) from bulk frontal Ctx and whole Str from CON (grey) and MIA (red) offspring indicating the sex [male (blue) and female (pink)] of each animal. n are individual animals. Data are shown as median ± interquartile range and analyzed by a linear mixed effect modeling and three-way ANOVA * p<0.05. **e,** IL-6 protein expression in maternal serum 3 hours after PIC injection. n are individual animals. Data are shown as mean ± s.e.m and analyzed using a two-way ANOVA with Tukey posthoc test. *p<0.05. Offspring from dams with a serum response below 1000 pg/ml were excluded from subsequent experiments.

